# Post-attack bioluminescence is an aposematic signal in cypridinid ostracods

**DOI:** 10.64898/2026.06.22.733700

**Authors:** Nicholai M. Hensley, Leanne M. Shulman, Trevor J. Rivers, Gretchen A. Gerrish, James E. Herbert-Read, James G. Morin

**Author notes:** **Corresponding author:** Nicholai M. Hensley. Department of Zoology, University of Cambridge, Cambridge, Cambridgeshire, United Kingdom (Nicholai M. Hensley).

## Abstract

Aposematic signals convey information about prey unpalatability to potential predators. Such signals are defined as advertisements, with prey usually signalling unpalatability before an attack. Overlooked, however, is whether signals presented after an attack can function as aposematic warnings and improve survival. Here, we test if post-attack signals can be aposematic by observing fish predators responding to prey (Ostracoda, Cypridinidae) that produce bioluminescence following an attack. By manipulating potential chemical defences of prey, and by comparing feeding responses to both bioluminescent and nonluminescent prey, we provide multiple lines of evidence to show bioluminescent prey are unpalatable and use post-attack bioluminescence as an aposematic signal. First, live, bioluminescent prey secrete bioluminescence only after being attacked. Second, predatory fishes rarely consumed bioluminescent prey, especially compared to nonluminescent alternatives. Third, because fishes readily ate bioluminescent prey that had been treated (frozen or boiled), which should remove chemical defences, bioluminescent species likely possess some unidentified defence over nonluminescent relatives. Finally, over the course of four experimental trials, predators were less likely to consume bioluminescent prey as their cumulative exposure to bioluminescent displays increased, indicative of learning. Post-attack aposematic signals, as found with bioluminescence, may be as common and effective a defensive strategy as pre-attack signals.

## Introduction

Many species use warning signals, like deimatic displays [1–3] or aposematism [4–6], to ward off attacks from predators. Classically with aposematism, prey advertise their toxicity or unprofitability before being attacked. Such visual signals must be conspicuous to predators, through colouration or contrast, and prey should be defended (i.e. chemically, physically, or behaviourally) to deter predators. Together, predators learn to associate these signals with defences and therefore reduce targeting defended prey. Aposematic colouration is well documented across terrestrial taxa [7–10], but rarely in the ocean (but see [11,12]).

While colour [4], salience [5], reliability [13], honesty [7], and frequency-dependence [14–17] can explain the origins and maintenance of aposematic signals, there has been less focus on whether the timing of aposematic signals affects their efficacy. Indeed, temporal variation in signalling may reduce the consistency with which information is conveyed, potentially weakening signal reliability and efficacy. Aposematic signals, therefore, are generally thought to be more effective if displayed prior to an attack [18]. Nevertheless, some species appear to display potentially aposematic signals after attack, which have been termed “hidden signals” [19], “facultative” [20], “switchable aposematism” [21], or “post-attack displays” [22,23]. In some cases, this dynamic signalling may be advantageous over static displays, for example, by balancing trade-offs between conspicuousness and crypsis [24]. But generally whether signals displayed after an attack can serve as aposematic signals and act to improve defence remains largely untested.

Biolumescence is one of the most common visual signals [25], mostly used by marine organisms [25–27]. Evidence that bioluminescence acts as an aposematic signal primarily comes from terrestrial systems that use bioluminescence as constitutive signals [28–31] (but see [32]). In marine systems, however, facultative bioluminescence seems the norm [27,33–37] with little evidence for an aposematic function [36] (but see [38,39]). In ostracod crustaceans (Tribe Lumini), for example, bioluminescence is facultative [35,40], and was co-opted from defence for mating [41]. Bioluminescence is produced when ostracods jointly secrete bioluminescent compounds [42,43], primarily an enzyme CLuciferase [44] (also called c-luciferase) and its high energy substrate vargulin [45], that react to generate blue light [44] (Fig. 1A).

**Figure 1.**
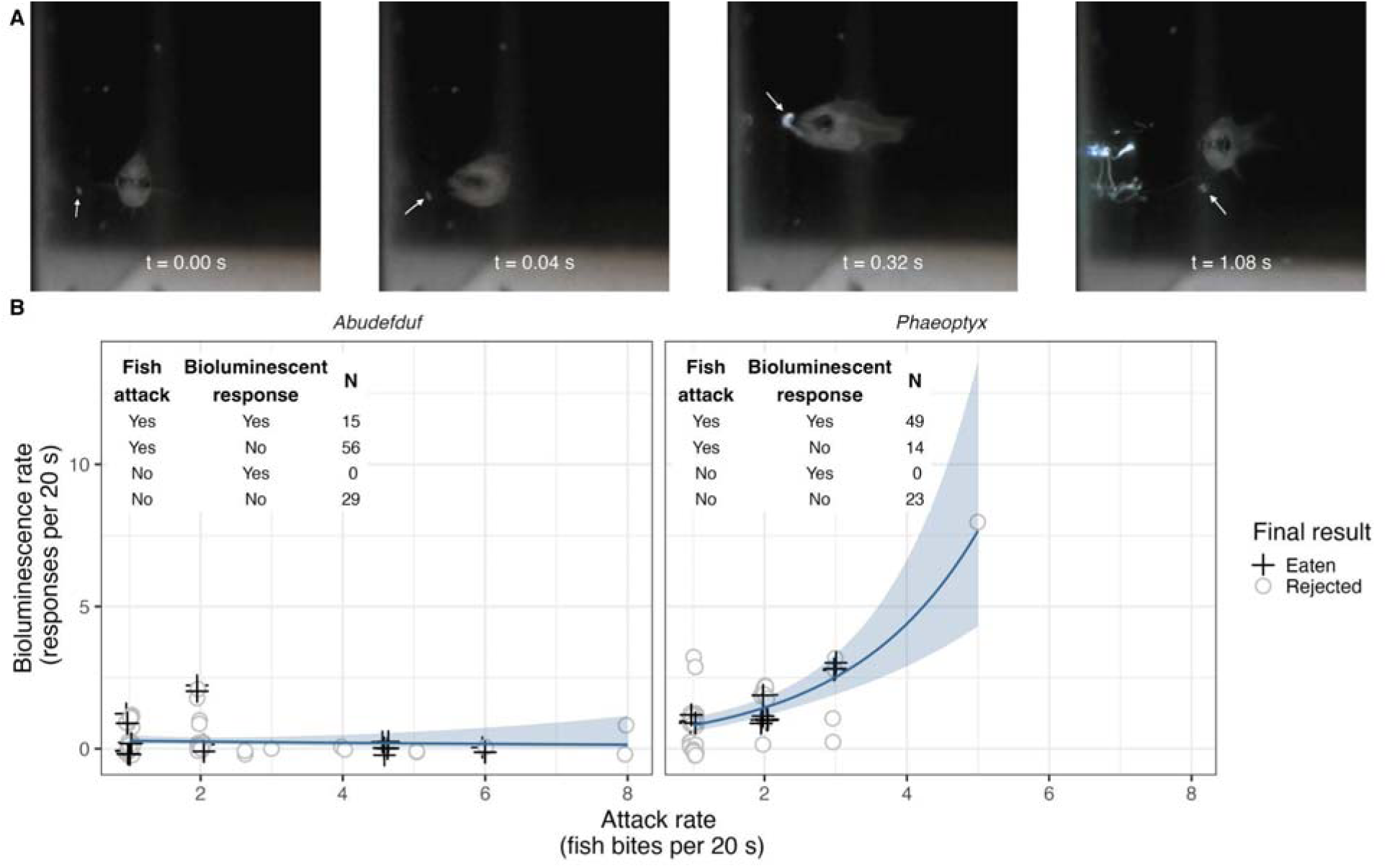
(A) Video frames showing a *Phaeoptyx* cardinal fish (i) targeting and (ii) attacking an ostracod, (iii) rejecting it as the ostracod releases bioluminescence (blob), and as (iv) the ostracod escapes leaving behind a glowing cloud. Ostracod indicated by arrows. Filmed under low red light. For full segment, see <u>here</u>. [51] (B) The rate of bioluminescent responses by living ostracods increased with *Phaeoptyx* attack rates, but were generally low for *Abudefduf* predators. Jittered points indicate whether food was eaten (crosses) or rejected (circles) in any *t* = 1 - 4. Insets: the number of time points when fishes attacked at least once, and the number of times points with at least one bioluminescent event.

Here we test if post-attack bioluminescence acts as an aposematic signal in ostracods. If so, we predict the following four key results. First, bioluminescent prey should only produce bioluminescence post-attack. Second, because bioluminescent prey should be defended, they should be consumed less compared to undefended, nonluminescent prey. Third, removing such defences should make bioluminescent prey more palatable than their untreated counterparts. And fourth, predators should learn that bioluminescent prey are defended when exposed to bioluminescent signals and decrease their likelihood to consume them.

## Materials and Methods

### Fish predators

All sampling and experiments took place at South Water Caye, Belize (16°48’46.8“N 88°04’57.3“W) from 15 - 23 March, 2012. We sought ecologically relevant predators (small-bodied invertivores), and to test for the generality of effects, selected both nocturnal and diurnal species. The species chosen represented those available in sufficient numbers. Nocturnal fish were cardinal fishes (*Phaeoptyx conklini* or *P. pigmentaria:* SL mean = 26.8 mm, range 15 - 40 mm; n = 8) collected above patch reefs one to two hours after sunset via mesh hand nets while snorkelling. For diurnal predators, we collected juvenile sergeant majors (*Abudefduf saxatilis*: SL mean = 19.5 mm, range = 13 - 26 mm; n = 11) using hand nets in < 50 cm water.

Fishes were immediately transferred to 2 L Ziplock bags with sea water and transported back to the lab, each housed in a 700 mL container (GladWare®) with ∼500 mL of fresh seawater and half a Toby Teaboy® tea strainer as shelter. Containers sat on shelves in an open-air lab at ambient conditions (∼ 12 L : 12 D, 27-30°C), shielded from direct sun with mesh nets as covers.

### Prey types and treatments

We provided predators with two different experimental prey types, each with four different treatments, to test whether bioluminescent prey were aposematic. The first prey type was bioluminescent ostracods (*Photeros annecohenae* Torres and Morin, 2007) [46], which we hypothesized were chemically defended, and can use bioluminescence when attacked. Our second prey type was nonluminescent ostracods (*Skogsbergia* sp. Poulsen, 1962) [47], which are ecologically similar to *P. annecohenae* but cannot produce bioluminescence and we suspect lack similar chemical defenses. For both species, only adult female ostracods were used as prey, and were readily caught using baited traps [48].

For both bioluminescent and nonluminescent prey, we used live, active ostracods to measure their natural palatability (hereafter “Live”). Second, we anaesthetised living ostracods in ∼2 : 1 0.36 M MgCl_2_ : sea water for > 30 min. (“Anaesthetised”). These animals retained physical and chemical defences like live animals, but could not produce bioluminescence following an attack, allowing us to disentangle behavioural and chemical defenses. Third, we froze ostracods in a drop of seawater for > 60 min. (“Frozen / Dead”). Lastly, we heat-killed ostracods in > 95°C sea water for 5 - 7 min. to denature bioluminescent or other defensive chemicals (hereafter “Boiled / Dead”), which at minimum would remove the ability to produce bioluminescence. The vargulin substrate is thermostable but oxidizes quickly [49], and the Cluciferase enzyme denatures under high heat. Treated prey were rinsed thoroughly in and transferred to ambient sea water before experiments. We also gave predators a third prey type as a palatable control (either raw fish muscle or a boiled cirolanid isopod) to test for general feeding motivation. Details on selecting prey can be found in the supplement and Fig. S2.

### Trial protocol

During a single experimental trial, each fish received one of 10 prey presentations at a time in a randomized order (2 control, 8 experimental prey). We used a 1 mL pipette to release prey just below the water’s surface with minimal liquid transfer (Fig. S1). We observed predator responses for 1 min. after the initial prey presentation, recording behaviors in 20 s intervals (hereafter time points; *t* = 1, 2, 3). Behaviors recorded were: (1) number of attacks: number of times prey was taken into the mouth or bitten; (2) number of rejections: if and when prey was expelled from the mouth; and (3) the number of bioluminescent events (for bioluminescent prey only). If prey was swimming (for live foods) or not being attacked at 20 s or 40 s, we used the pipette to capture and re-release it near the water’s surface, making it more visible while sinking. If prey was eaten, we marked it during that time point. If the prey was uneaten after 1 min., we left it with the fish for ∼8 - 11 min. while we tested the remaining fish. After all fish had received a prey item and been observed for 1 min., we went back and marked the final result (eaten or uneaten) of the prey for each fish (at *t* = 4, usually 8 - 11 min. later; Fig. S1). If the prey was uneaten, this was recorded and the prey removed. If the prey was eaten between *t* = 3 and *t* = 4 while we were watching different fish, we added 1 to the total attack number for the focal fish with that prey, inferring that prey must have been attacked at least once to be eaten. After collecting the data for the first prey presentation for each fish, we would then start the next series of presentations, repeating the process until all 10 prey had been given to every fish and completing the trial. At the end of each trial, we replaced the water in the container with fresh seawater. Each fish had 4 trials over 5 days. Trials occurred at ambient temperature (27 - 30°C), and were run separately for nocturnal predators (between 1900 and 2300 hrs) and diurnal predators (between 1000 and 1700 hrs). All fish within a species were tested during a trial. For more details, see the supplementary information and Fig. S1.

### Statistical analysis

We excluded three trials (1 *Abudefduf* and 2 *Phaeoptyx* trials out of 78 total) because none of the 10 prey were attacked within that trial (Fig. S3). We only used data from prey that were attacked at least once during *t* = 1 - 3. We did not include control prey in our analyses as these were to only assess motivation, but present them in Fig. S4. For a small number of prey presentations (n = 9), we could only record the total number of attacks or bioluminescent events in 1 min., not in 20 s intervals due to high attack rates. To include these data conservatively, we divided the total number of attacks or bioluminescent responses evenly across *t* = 1 - 3, which should bias our rates against our expected results without removing the trial completely. Some *Phaeopteryx* predators received ostracod prey during their pre-trials (n = 5); we included these in subsequent analyses to account for their potentially increased experience. We combined Frozen and Boiled into a single treatment level (“Dead”) as there was no statistical difference in the proportion eaten between these treatments (Fig. S5).

We analyzed the remaining and recategorized dataset in R (vers. 4.4.0) with RStudio (vers. 2024.04.2+764) to address the following questions:

### Are bioluminescent events and predator attacks related?

We used a Fischer’s Exact test to ask if the number of time points with and without fish attacks is correlated with the number of time points with at least one or without any bioluminescent events. We limited our analysis to presentations with live bioluminescent prey and between *t* = 1 - 3. This method cannot account for repeated measures, but is exact with imbalanced contingency tables, as we observed (Fig. 1B, insets).

With the same subset of data, we also tested if the number of attacks predicted the number of bioluminescent events. To do this, we fit a generalized linear mixed-effects model with a Poisson error distribution to test whether the number of bioluminescent events was influenced by the number of attacks, predator genus, and their interaction. Time point and presentation order were included as additional fixed effects, and fish identity was included as a random intercept to account for repeated observations of the same individual. The number of attacks was continuous, predator genus was categorical, and both time point and order of presentations were ordinal.

### Are bioluminescent prey unpalatable?

To assess prey palatability, we fit a discrete-time hazard mixed-effects model to our time-to-event data (Fig. S6). This approach allowed us to test whether prey type and treatment affected the probability of prey being eaten during each time point, conditional on it having remained uneaten until that time point [50]. Prey type and treatment were fitted as categorical fixed effects. Preliminary plots of the proportion of prey eaten (calculated as the number of presentations in which the predator consumed the offered prey at any time point, divided by the total number of presentations) showed consistent treatment effects across prey types, so their interaction was not included (Figs. S4, S5). Time point within presentation (*t =* 1 - 4) was included as an additional fixed effect, and nested random effects including fish identity accounted for repeated measurements of the same predator. A discrete-time approach was used because the observation intervals differed in duration: the first three intervals each lasted 20 s, whereas the final interval lasted 8 - 11 min. Statistical comparisons were performed using the relative risks estimated by the model, but results are presented as cumulative probabilities of being eaten over all time points, derived from the corresponding survival function, to aid interpretation. Further statistical details are provided in the supplementary information and Fig. S7.

### Do predators modify their behaviour with experience of bioluminescent events?

To test if consumption risk changed with exposure to bioluminescent events, we summed each fish’s cumulative exposure to bioluminescent events over all time points, prey presentations, and across trials as a continuous variable called bioluminescence experience. We added this bioluminescence experience variable as a predictor to the same discrete-time hazard model described above, also including trial number as a continuous variable because experience was potentially correlated with repeated testing. We included interactions between prey type and bioluminescence experience, and between prey treatment and trial number, to test whether the effect of experience differed among prey types, and whether treatment effects changed over successive trials. We centered continuous variables in models, and plotted marginal means for significant terms on the original scale.

During reduction of the global model using the “drop1” function, we removed the following predictors: predator genus, and the order of prey presentation (Likelihood ratio tests; p = 0.7837 and p = 0.3541, respectively). Interaction terms between prey type and trial number, and between prey treatment and bioluminescence experience were not included in the most explanatory model (Likelihood ratio tests; p = 0.3259, p = 0.4319, respectively). For details of the global model, variable reduction, random effects structure, and final model selection, see the supplementary information.

We used the function “glmmTMB” in the package ‘glmmTMB’ to build all mixed effect models. We performed post-hoc pairwise comparisons using the ‘emmeans’ package, with the function “emmeans” for comparisons of effect means for categorical variables specifying a Sidak correction for multiple comparisons, and “emtrends” for comparisons of effect slopes for continuous variables.

## Results

### Bioluminescent events occurred after, and increased with, nocturnal predator attacks

Bioluminescent events with live, bioluminescent prey never occurred without at least one attack (Fig. 1B, insets), and never before one (Fisher’s exact test, p < 0.0001). The rate of bioluminescent events increased with the rate of predator attacks for nocturnal predators (β_Number of attacks x Genus_ *_Phaeoptyx_* = 0.65, 95% CI [0.27, 1.03], p = 0.001; Fig. 1B, Table S1) but not diurnal predators (β_Number of attacks_ = −0.09, 95% CI [−0.43, 0.24], p = 0.574; β_Genus_ *_Phaeoptyx_* = 0.44, 95% CI [−0.43, 1.32], p = 0.322; see the supplementary materials for a discussion). This effect was unchanged when a single outlier was removed (Fig. S8; Table S2).

### Bioluminescent prey were eaten less than nonluminescent prey

Prey type had a significant effect on the risk of consumption (β_Nonluminescent_ = 0.66, 95% CI [0.42, 0.91], p < 0.001; Table 1). Post-hoc pairwise comparisons confirmed that live nonluminescent prey, anaesthetised nonluminescent prey, and dead nonluminescent prey were more readily eaten than their bioluminescent counterparts (Fig. 2; Table S3).

**Figure 2.**
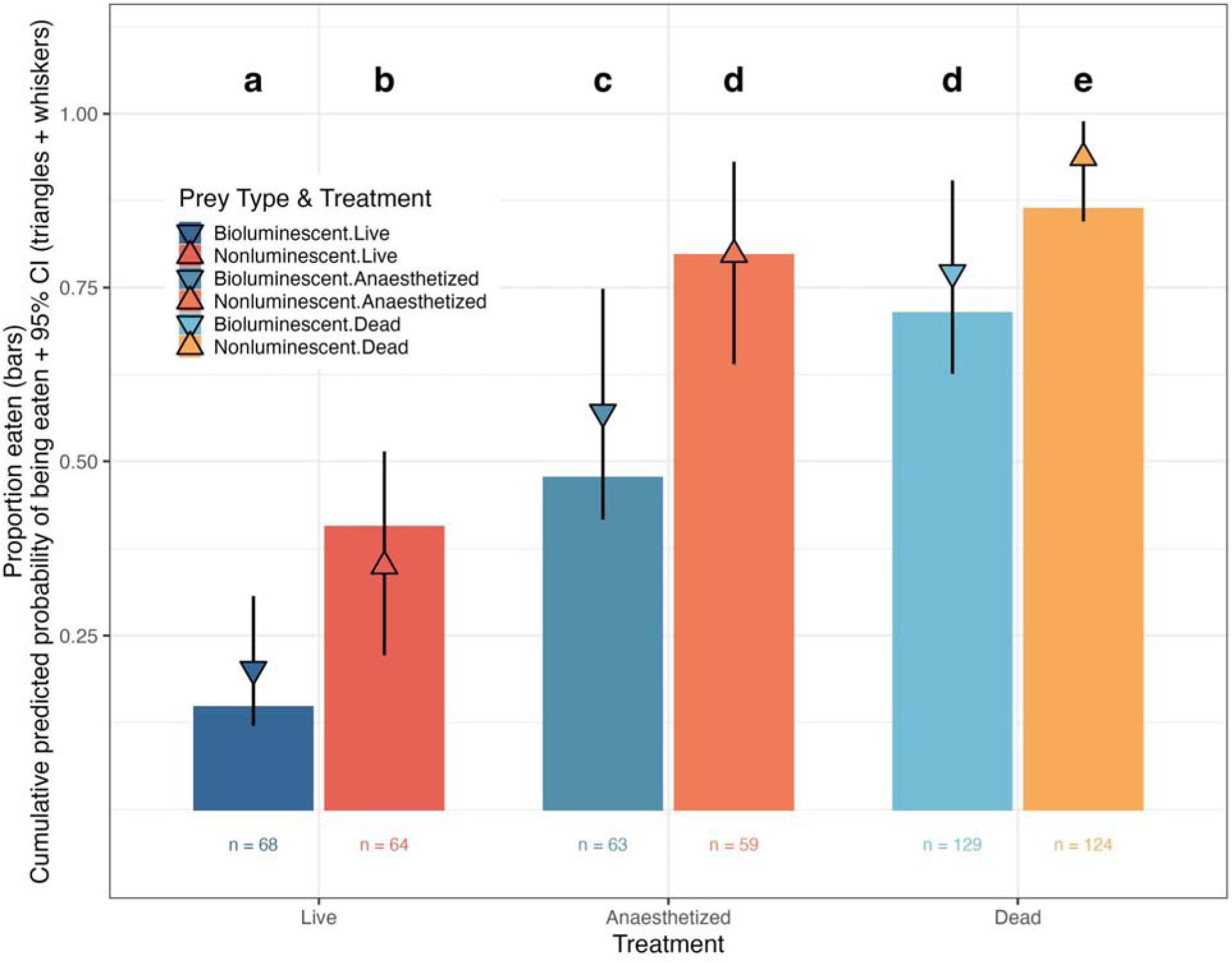
Proportion of prey eaten (y-axis) across all trials for each prey type (colours) and treatment (x-axis). Bars are the proportion of feeding presentations when eaten. Triangles are the marginal cumulative probabilities of being eaten from the discrete-time hazard model for type (blue, downward triangles = bioluminescent; red, upward triangles = nonluminescent) and treatment (darkest = live; brightest = dead). Whiskers are 95% bootstrapped confidence intervals. Letters denote different groups based on non-overlapping bootstrap distributions.

**Table 1.** Results from the best discrete-time hazard model on the risk of being eaten as a function of the prey type, treatment, the cumulative experience of each predator with bioluminescent events, and over consecutive trials, along with interaction terms and random effects.

### Live prey were eaten less than treated prey

Treatment had a significant effect on the risk of consumption (β_Anaesthetised_ = 1.36, 95% CI [0.94, 1.77], p < 0.001; β_Dead_ = 1.92, 95% CI [1.54, 2.31], p < 0.001; Fig. 2, Table 1). Post-hoc pairwise comparisons confirmed that anaesthetised and dead prey were more at risk of being eaten than live prey for both bioluminescent and nonluminescent types (Fig 2; Table S3). Dead prey were also more likely to be eaten than anaesthetised prey for both bioluminescent and nonluminescent types (Fig 2; Table S3).

### Bioluminescence experience reduced the risk of being eaten for bioluminescent prey only

Experience with bioluminescent responses during attack significantly decreased the risk of being eaten (β_Luminescence experience_ = −0.12, 95% CI [−0.19, −0.05], p < 0.001; Fig. 3; Table 1). However this differed between prey types (β_Nonluminescent * Luminescence experience_= 0.10, 95% CI [0.05, 0.16], p < 0.001; Table 1). Post-hoc pairwise comparisons confirmed that for nonluminescent prey, predation risk was and remained relatively high despite increasing experience with bioluminescent responses. In contrast, increased exposure to bioluminescent responses was associated with a decline in the risk of consumption for bioluminescent prey (Table S3). This translated into different cumulative probabilities of being eaten for bioluminescent versus nonluminescent prey as a function of predator exposure to bioluminescent responses (Fig. 3).

**Figure 3.**
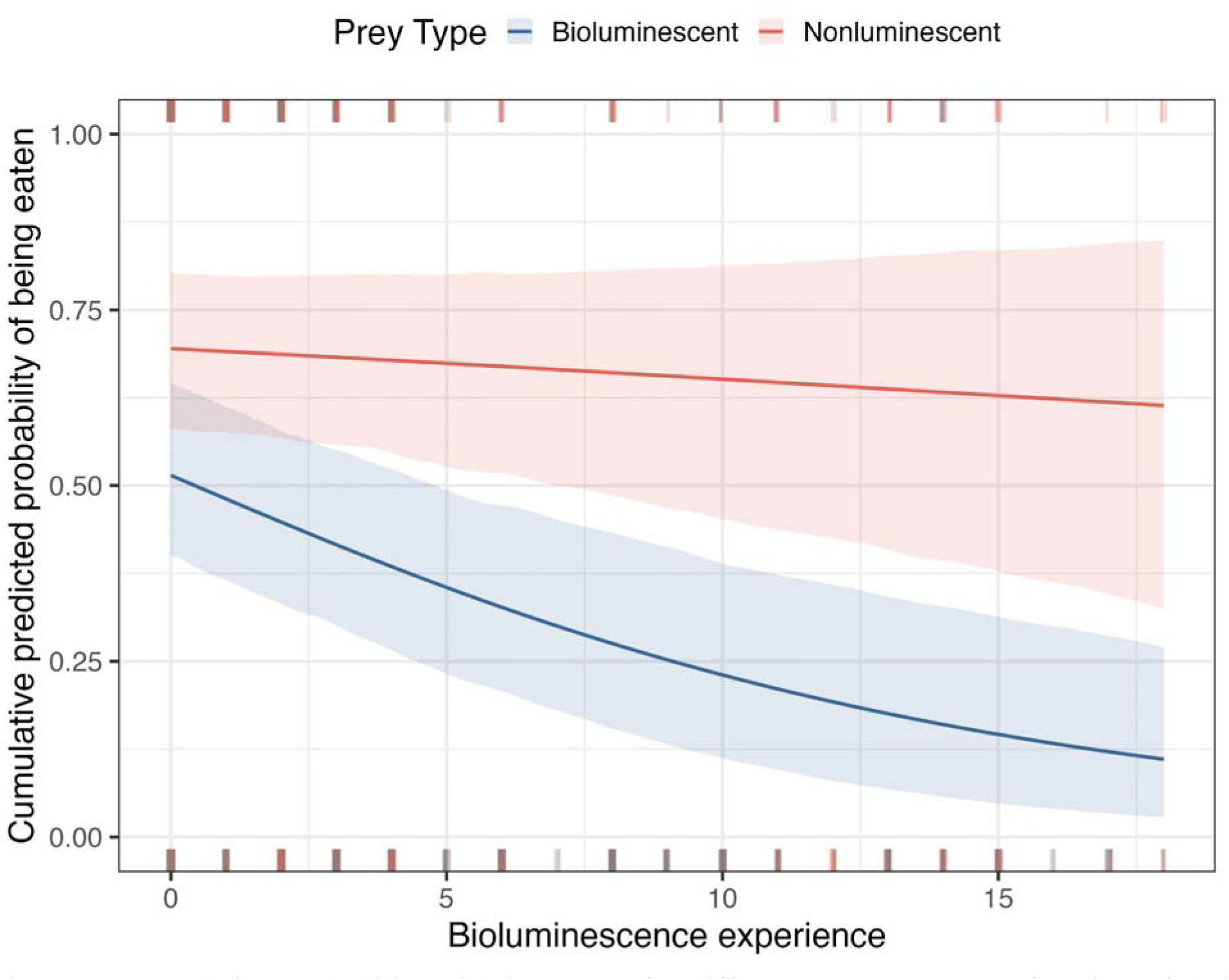
Cumulative probability of being eaten for different prey types as a function of their cumulative exposure to bioluminescent responses. As experience increased (x-axis), cumulative probability of consumption (y-axis) decreased for bioluminescent prey. Model predictions are marginal cumulative probabilities with all other fixed effects held at their mean and plotted on the original scale. Shaded regions are 95% confidence intervals from bootstrapping. Y-axis is the cumulative probability of being eaten (0 = no cumulative probability of being eaten, 1 = 100% cumulative probability of being eaten). Jittered raw data (0 = uneaten, 1 = eaten) as ticks coloured along x-axes.

## Discussion

Here, we show bioluminescence can be a post-attack signal to teach predators about prey unprofitability using four lines of evidence. First, ostracods produced bioluminescent signals only after being attacked. Second, nonluminescent prey were more readily eaten than bioluminescent prey, regardless of treatment. Third, when prey were dead or treated to remove their potential defences, consumption of bioluminescent prey increased. Last, predatory fishes reduced their likelihood of eating bioluminescent prey as their exposure to post-attack, bioluminescent responses increased, an indication of predator learning.

Our results suggest aposematic bioluminescence is maintained because such defensive signals are likely honest. Bioluminescent prey were consumed less than non-bioluminescent prey even when anaesthetised, suggesting differences in consumption were not due to differences in escape or behavioural defence between prey species. Anaesthetised bioluminescent prey were also consumed less than dead bioluminescent prey that were boiled or frozen. Boiling, and even freezing, is likely to destroy proteins or small molecules that may act to increase unpalatability. Indeed, recent work suggests ostracods may possess toxin-like genes expressed in their bioluminescent secretion organ [52].

We also found that exposure to light emission after an attack appears to teach predators to reject defended prey. This is surprising because constitutive defensive signals (e.g. static colouration) are more common in nature and reliably teach predators about defense [18]. Moreover, post-attack signals are theoretically less effective for the associative learning that maintains aposematism [53]. However, certain scenarios may favour post-attack facultative signalling. First, predators can learn to avoid prey with post-attack signals (i.e. backward conditioning) [22]. Second, prey may need to balance trade-offs between signalling energetics [40] and the benefits of inconspicuousness. And third, if predator learning about defended prey is slow, highly variable, or not maintained, which increases the relative cost of high signalling [21]. Because marine bioluminescence is largely facultative and restricted in colour [54], post-attack bioluminescence may be a more common aposematic strategy than appreciated.

A key question is how post-attack aposematism could initially evolve. First, bioluminescence could have been co-opted from other defensive functions [27]. The “burglar alarm hypothesis” suggests that bioluminescence could be an alarm call attracting primary to secondary predators [55], shown in dinoflagellates [39,56,57]. Bioluminescence could also startle or distract predators (i.e. deimatism, “smokescreen” hypotheses), increasing prey escape [27,36,56]. If defensive bioluminescence arose via other such functions, secondary defensive traits (e.g. toxins) could subsequently evolve and allow signal co-option into aposematism. Firefly bioluminescence evolved before specific toxins (although not necessarily toxicity) [58], suggesting co-option is plausible. Second, bioluminescent prey like ostracods often survive attack [38,59], which may relax the assumptions required to evolve aposematism. Theory states that defended prey with conspicuous, novel signals and who are preferentially targeted by naive predators can only multiply if their inconspicuous kin pass along defensive traits until predators learn to avoid the signal [60–62]. Instead, bioluminescent defended prey may survive attack to pass along both signal and defensive traits, even as naive predators learn to avoid them.

Overall, our findings support defensive bioluminescence in ostracods is a post-attack aposematic signal that enables predators to learn about prey unprofitability. By remaining inconspicuous until attacked, prey may gain the benefits of defensive signals while limiting costs of continuous conspicuousness. Co-option from other defensive functions, together with the capacity to survive predator attacks, could help explain the evolutionary origins of this strategy. These results broaden our understanding of aposematism by showing that warning signals can be effective even when expressed only after an attack has begun.

## Author Contributions

NMH curated, managed, and analysed data. LMS collected data and advised on analysis. TJR, GAG designed experiments and collected preliminary data. JHR contributed to data analysis. JGM conceived the project, designed experiments, and collected and analysed data. NMH wrote, and TJR, GAG, JHR, LMS, and JGM edited and approved of the manuscript.

## Acknowledgements

Thanks to EK Bladon for advice on statistical modeling and comments from JJ Arbon on a draft of this work.

## Ethics Statement

Data were collected at Southwater Caye, Belize under permit 2012 (21) VOL IX. IACUC approval of protocols was obtained from UW La Crosse under protocol #4-15.

## Data availability

Data will be made available on the corresponding author’s GitHub, along with analysis scripts, here: https://github.com/NikoHensley/Aposematic_defenses

## Supplementary Information

### Supplementary Methods

#### Fish predators

As an alternative diurnal predator, we also collected damselfishes using the same methods described in the main text (n = 9; *Stegastes adustus* and *S. leucostictus*, SL = unmeasured). Two damselfishes were immediately released after refusing any offered food. The remaining 7 fish were used in pre-trials and into at least one experimental trial. During trials, we observed that damselfish did not readily eat, and if they did, their attacks did not produce bioluminescent responses. We subsequently released most *Stegastes* after a single trial; any data we collected were excluded from analyses.

#### Ostracod prey

Both ostracod species are nocturnal, belong to the same family (Cypridinidae), and are scavengers that come to the same baited traps with the same bait at the same time in the same habitat. They are roughly the same size (∼ 1.25 - 2.0 mm adults, depending on sex), and share a general ostracod bauplan consisting of a bivalved shelled carapace that encapsulates their entire body, which has 8 pairs of limbs, reduced body segmentation relative to other crustaceans, and two large lateral eyes. These species are ∼20 Mya diverged from one another and possess finer morphological differences that distinguish them; they may possess other differences in biochemical defenses which are unmeasured here.

#### Trial protocol

Before experimental trials, we did three preliminary tests with potential, common foods, (mysids, cumaceans, cirolanid isopods, and nereid polychaetes) and one pre-trial with four prey types, one of which was the control food. Based on the results of these preliminary tests, we chose the cirolanid isopod as an appropriate control food as it was eaten in a relatively high proportion of trials where fish ate (Fig. S2), and was more easily obtained from the wild. For the first experimental trials, we used raw fish muscle as the control food before switching to isopods, which required time to collect en masse.

For trials at night, we used red head lamps to visualize containers and behaviors. After preliminary tests and pre-trials, fish only received food during experimental trials and were kept without food for two days prior to starting. Two researchers performed the experiments working as a pair: one prepared the food, while another performed the observations. All experiments were manually coded without the use of video.

For coded behaviors, we could only directly observe occurrences during time point *t* = 1 - 3 due to our sampling design (Fig. S1). During these time points, the number of attacks and number of bioluminescent responses were directly observed. However, fish were unobserved between time points *t* = 3 and *t* = 4, as mentioned in the main text. This prevented us from directly observing any bioluminescent responses for this time point. For the number of attacks at *t* = 4, we inferred if at least one attack occurred if prey was eaten. Both uneaten and eaten prey may have been attacked more than this between *t* = 3 and *t* = 4, but we cannot account for this variation, and do not attempt to do so in our analyses.

#### Statistical analyses

##### The discrete-time hazard model

To assess if both prey types, treatments, and predator experience influenced the risk of being eaten, we built a discrete-time hazard model, with various fixed effect predictors and random effects, aimed at addressing each factor jointly but that would take into account our sampling design and repeated measures. Discrete-time hazard models are a form of survival model, which provide predictions on the relative instantaneous risk of an event occurring (i.e. being eaten) given the probability that a sample has survived until that time [50]. Relative risks can be transformed into a cumulative probability of survival via a survival function that integrates the entire relative risk over the observation time (in our case, time points *t =* 1 - 4) (see below). The global model appeared as:

eaten ∼ time point + prey type + prey treatment + trial number + order + bioluminescence experience + predator genus + prey type:time point + prey type:trial number + prey type:bioluminescence experience + prey treatment:time point + prey treatment:trial number + prey treatment:bioluminescence experience + (1 | sample ID / trial ID / fish ID), family = binomial(link = “cloglog”)

We included a term for the order of prey presentations within a trial (“order”) as an ordinal variable. We observed that fish behavior changed over the course of trials, and so included two-way interactions between prey type and trial number, and prey treatment and trial number. Because we were interested in testing if fish changed behavior based on experience with different prey, we included two-way interactions between prey type and bioluminescence experience, and between prey treatment and luminescence experience in the model. We included a categorical variable for the genus of the fish predator. See below for an explanation of random effect variables.

During analysis, we centered continuous variables to reduce variance inflation factors below 5 for model selection of interaction terms. We used iterative rounds of the “drop1” function to determine predictor term importance, and retained terms or interactions that were in models with marginally or significantly better fit via Chi-square tests (p < 0.07). We used the “test_performance” and “compare_performance” functions from the ‘performance’ package to assess the best model with Bayes factors. We assessed multicollinearity of variables using “check_multicollinearity” and retained variables in the model that had adjusted VIF < 3 and tolerance > 0.1, preferentially removing interaction terms with high adjusted VIF if they could not be resolved.

Our final model after variable reduction was:

eaten ∼ time point + prey type + prey treatment + trial number + bioluminescence experience + prey type:bioluminescence experience + prey treatment:trial number + (1 | sample ID), family = binomial(link = “cloglog”)

##### Translating from relative risk to cumulative probability

Our discrete-time hazard model reports the relative risks of observing our focal behaviors, either being eaten or being attacked. In general, relative risks can be translated into cumulative risk by integrating the risk over the observation time. First, we transformed the linear relative risk (as estimated by the model) into a risk probability using the complementary log of logs, such that:

Risk probability = (1 - *e^-^e^instantaneous risk^*)

We then integrated these risk probabilities across *t* = 1 - 4 using their cumulative product from each time point. By subtracting this cumulative risk probability from 1, we generated a classic survival function from the discrete-time hazard model, which describes the cumulative probability of prey surviving. We again subtracted this cumulative survival probability from 1 to get the cumulative probability of *not* surviving (i.e. of being eaten / consumed). The statistics in our main text are presented on the linear relative risk, and tables present them as the risk ratios, but we graph the results on the cumulative probability of being eaten as it is conceptually easier to compare and interpret across model predictors. We present visualizations of how the risk probability is integrated over time in Fig. S7 and across relevant predictors. Figs. 2 and 3 in the main text, and Figs. S7 and S9 in the supplement all show results for the fixed effects on the marginal cumulative probability of being eaten.

To similarly translate the confidence in our marginal risks to their marginal cumulative probabilities, we used a bootstrapping method. For each model, we took the confidence intervals from the model and the covariance matrix between predictors and generated 1,000 coefficients per focal predictor, and across the range of values for the other categorical or ordinal predictors, and while holding all other continuous predictors at 0 (their average value after scaling). These 1,000 simulated marginal risks were then cumulatively multiplied across *t* = 1 - 4 to create confidence intervals for the marginal cumulative probabilities. To compare differences between categorical groups, as in Figs. 2, 3, S5, and S9, we looked at the pairwise differences between groups for these 1,000 simulated marginal cumulative probabilities. If 95% of the distribution of differences between any two groups overlapped zero, we considered them statistically indistinguishable.

##### Random effects structure for both linear and discrete-time hazard models

Both mixed effect models - the first testing the rate of bioluminescent responses as a function of the rate of attack, and the second testing the risk of consumption using the discrete-time hazard approach - included the same nested random effect structure initially because fishes were observed for multiple time points and multiple trials. We nested observations within a prey presentation (“Sample ID”) within a trial (“Feeding Trial”) within each fish (“Individual ID”). For each model, we removed random effects that explained little to no variance ( < 0.001) and that did not improve the dispersion of the model if retained. For each mixed effect model, we compared models with the same fixed effects (chosen after using “drop1”, as described above) but with either a reduced random effect structure, removing effects that explained little variation, or the complete nested, random effects structure using the “check_performance” function. We retained the model with a random effect structure that produced the lowest BIC.

##### Do predators modify their willingness to attack prey over time or with experience of bioluminescent events?

An important prediction of the aposematism hypothesis is that experienced predators will reduce targeting prey that honestly signal their defences. However, in our system signals only occur after an attack. Therefore, if predators do learn about prey defences from post-attack bioluminescent signals, they should readily attack prey because there is no advertisement signal, but still reduce their consumption as they are exposed and learn to associate the post-attack signal with defences. In addition, although our discrete-time hazard model can account for varying cohort size over the observation period, variation in attack risk may influence variation in the risk of consumption, potentially confounding our estimates of consumption risk. Therefore, we also modeled the risk of attack to different prey types and treatments over changes in predator experience with bioluminescent responses, and over time (number of trials) in a model, just as we did with the risk of being eaten.

To assess if prey type and prey treatment affected the risk of being attacked, we performed the same statistical analysis as we did for the risk of being eaten (above, and main text) but using the global model with the full time-to-event dataset for attacks (Fig. S10), but not filtering out whether food had been attacked at least once (Fig. S3A versus S3B). One round of model fitting indicated that interactions between time_point:prey_type and time_point:prey_treatment were too collinear to include as assessed by VIF > 3. After removing these terms, the final model retained was:

attacked ∼ time point + prey type + prey treatment + trial number + order + luminescence experience + predator genus + order:trial number + prey type:trial number + prey type:luminescence experience + prey treatment:trial number + prey treatment:luminescence experience + (1 | sample ID / trial ID / fish ID), family = binomial(link = “cloglog”)

##### Does bioluminescence experience, or exposure to defended prey, better explain the cumulative probability of being eaten?

To assess if exposure to bioluminescent events (i.e. bioluminescence experience) was a better predictor of the probability of predator consumption than simply exposure to a defended prey, we calculated a metric of exposure to bioluminescent ostracods as the cumulative sum of attacks to live bioluminescent prey, regardless of whether they produced bioluminescent displays or not. Within an individual predator and across all feeding presentations and trials with bioluminescent ostracods, we calculated the cumulative sum of attacks. Using our best fit discrete-time hazard model, we re-ran the analysis including this metric of exposure to defended prey, as well as their cumulative experience with bioluminescent responses. This model appeared as:

eaten ∼ time point + food type + food treatment + trial number + bioluminescence experience + defended prey attack experience + food type:bioluminescence experience + food treatment:trial number + (1 | fish ID), family = binomial(link = “cloglog”)

We then used the “drop1” function to decide which of the included fixed effects should remain in the model, as stated. We compared this model with the reduced model using Bayes Factors, AICc, RMSE, and marginal R^2^ values.

### Supplementary Results

#### Time in presentation or presentation order did not affect the number of bioluminescent events

The rate of bioluminescence responses by live prey within any presentation may be subject to the dynamics of predator attacks, which vary over time. To understand if time points within a presentation or if presentation order affected the number of bioluminescent events, we included these two predictors in our linear mixed effect model. When compared to a model with or without each of these single predictors, neither model with time point nor order were statistically better fit than without (Likelihood Ratio Test, p = 0.3719, p = 0.7323, respectively), and so these variables were excluded from the final model. The best fit model included the number of attacks and the genus of the predator, as presented in the main text.

#### The relationship between the presence and number of attacks on the presence and number of bioluminescent events is robust to outlier inclusion

To understand if the relationship between the number of bioluminescent events and the number of attacks per time point was sensitive to one high value, we excluded this trial from analysis. This had no meaningful effect on our results; the number of bioluminescent events was significantly predicted by the number of attacks for *Phaeoptyx* nocturnal predators (β_Number of attacks x Genus_ *_Phaeoptyx_* = 0.63, 95% CI [0.18, 1.07], p = 0.006; Fig. S8; Table S2).

#### Although the risk of being attacked varied over time at different timescales (within presentations, between presentations, and between trials), the cumulative probability of being attacked over trials and with increasing bioluminescence experience did not vary

Although a discrete-time hazard model is a statistical approach to handle changes in the size of a cohort as events occur and prey drop out of the observation period (i.e. from being eaten), we wanted to quantify how the cumulative probability of being attacked may influence our understanding of the cumulative probability of being eaten, as they are sequentially dependent. To do so, we used a discrete-time hazard model for the risk of being attacked, integrating over the time periods (*t* = 1 - 4) and converting the relative probability of not being attacked into the cumulative probability of being attacked using the product of the survival function (as described above for discrete-time hazard models).

Although the instantaneous risk of being attacked varied by a number of intrinsic, experience-dependent, and time-dependent factors as we report in Table S4, these risks did not translate into meaningful differences in the cumulative probability of attack (Fig. S11, S12). There are some trends in the cumulative probability of being attacked, such as nonluminescent prey being marginally less likely to be attacked than bioluminescent prey (Fig. S11A). This seemed to differ over successive trials, where nonluminescent prey initially had a low predicted cumulative probability of being attacked for predators with higher experience of bioluminescent responses, but this was never significantly different from the predicted probability of being attacked for bioluminescent prey (Fig. S11B). Differences in these trends only decreased with increasing trials, regardless of experience with bioluminescent responses. Similarly, different prey treatments had trending differences in their marginal cumulative probabilities of being attacked, but these differences were never significant and became less extreme with more time and more experience (Fig. S11C).

Overall, the cumulative probability of being attacked was generally high and never differed significantly over time, with experience, or by treatment. Although we see a slightly higher probability of attack for bioluminescent prey over nonluminescent prey, this relationship would work counter to our results on the cumulative probability of being eaten, which is consistently lower for bioluminescent prey. These results support our general conclusions that predator attack rates remained high throughout trials, sufficiently to reliably estimate the cumulative probability of consumption, and despite the fact that predators experienced post-attack defensive displays.

#### Outside bioluminescence experience, prey treatments differed from one another and over subsequent trials in their risk of being eaten regardless of prey type

Predators may change their willingness to eat different prey over time, simply as a function of captivity or dynamic internal states (i.e. hunger). As time was potentially correlated with bioluminescence experience, we included it in our discrete-time hazard model as mentioned above to jointly estimate the effects of learning with the effects of time.

From our discrete-time hazard model on the probability of being eaten, we find that the number of trials significantly impacted the risk of being eaten (β_Trial number_= −0.31 95% CI [−0.63, −0.00747], p = 0.045; Fig. S9; Table 1), however this differed for the three treatments (β_Anaesthetised * Trial number_ = 0.41, 95% CI [0.05, 0.78], p = 0.027; β_Dead * Trial number_ = 0.61, 95% CI [0.27, 0.94], p < 0.001). Both anaesthetised and dead prey had a higher risk of being eaten compared to live prey (Table 1), with live prey having a lower cumulative probability of being eaten across trials (Fig. S9). Post hoc comparisons indicated a slight increase in the risk of anaesthetised prey being eaten across successive trials, however, this trend did not differ significantly from zero or from those observed for dead or live prey (Table S3). These effects were estimated jointly with the effect of bioluminescence experience, indicating that despite changes in the risk of consumption over time for different prey treatments, bioluminescence experience still explained a significant portion of variation in predation risk.

#### Models with a metric of exposure to defended prey are no better at explaining variation in the risk of prey consumption than the metric of bioluminescent signalling

When including the cumulative number of attacks on defended prey as a metric of predator exposure, we find that models with this variable are no more preferred than without (Wald Type II χ^2^ test, p = 0.1868). When comparing models that have either defended prey experience or bioluminescence experience, we find that the model with the bioluminescence experience has a lower corrected AIC (1117.9 versus 1132.2), a higher marginal R^2^ (0.377 versus 0.347), and a slightly lower root mean squared error (RMSE 0.355 versus 0.358). Comparing the models directly using Bayes Factors shows marginally greater support for the model with the cumulative attack experience (BF < 0.001). Together these analyses support that the model with bioluminescence experience is largely preferred than the model with attack experience.

### Supplementary Discussion

Our results indicate variation across predator genera in the ability to invoke a bioluminescent response from bioluminescent prey (Fig. 1); cardinalfishes received the most bioluminescent responses despite prey having a higher relative risk of attack in trials with sergeant majors (Figs. S11, S12; Table S4). It is tempting to speculate on the nature of this variation, as fishes differ in their ecology: sergeant majors are diurnal invertivores, whereas cardinalfishes are nocturnal, and when we might expect bioluminescent signalling to be most effective. However, these species diverged millions of years ago one another, are in different families, and differ in many ways besides foraging niche (size, habitat, etc.). To test if predator foraging ecology is related to the anti-predator response of bioluminescent prey, one might take a comparative approach testing the same prey and their probability of producing bioluminescent signals when paired against many different predators that vary in their foraging ecology (i.e., diurnal versus nocturnal), and which have been sampled in a phylogenetically informed way. Our data present baseline information for two such species, but lack power to test that question explicitly.

An alternative explanation for the lack of bioluminescent responses by bioluminescent ostracods to diurnal predators is that the ostracods tested during the day were asleep. Recent work shows that a congeneric species shows physiological hallmarks of a quiescent state during the day [63], and that these animals are less reactive when perturbed by movement (such as water from a pipette, pers. obs.). However, ostracods can be awake during the day, as animals will swim around, feed, and even produce bioluminescent responses if disturbed (pers. obs.; amongst *Abudefduf* predators in Fig. 1). Exploring factors explaining variation in prey response is an apt future goal.

---

**Figure S1.**
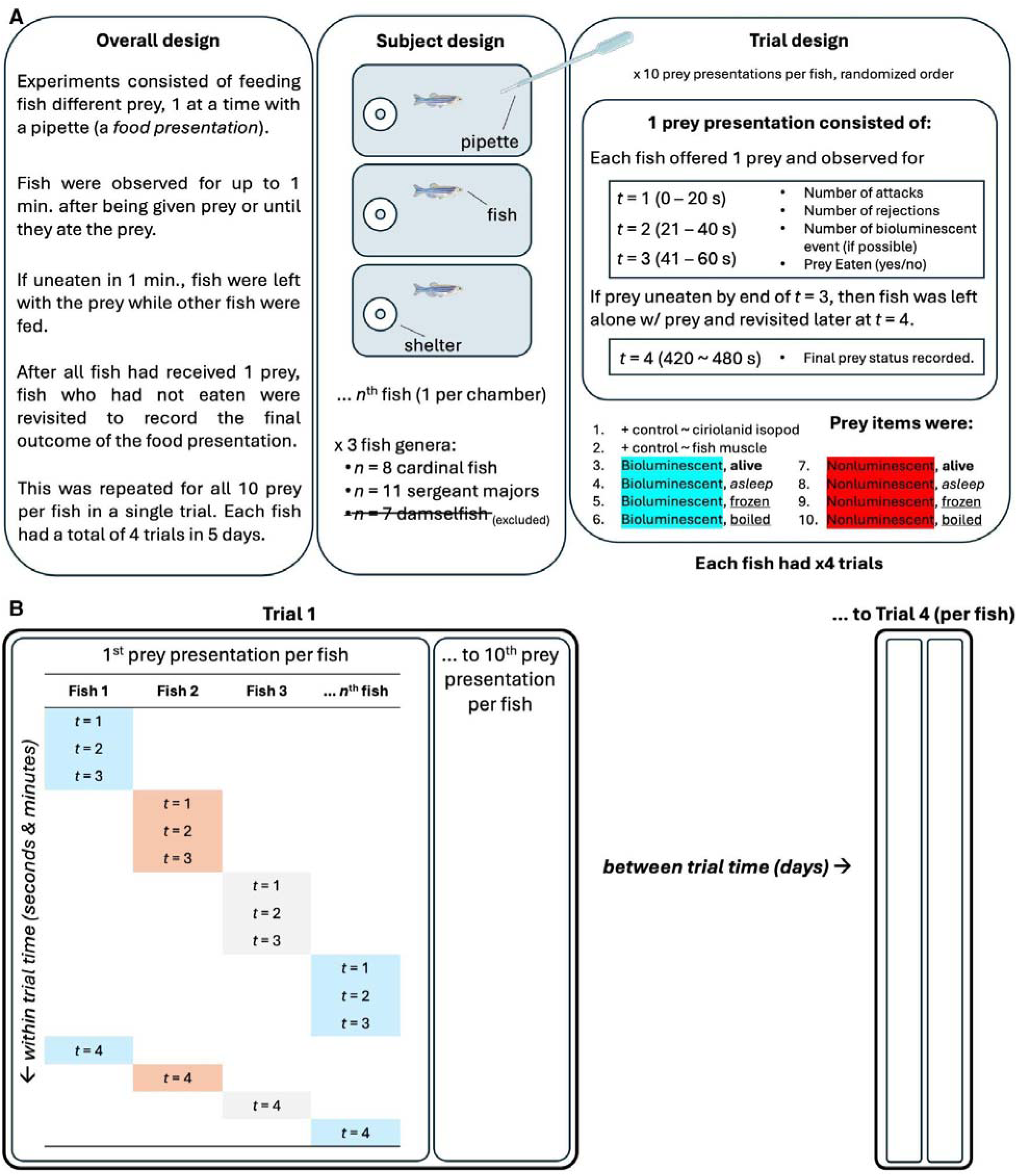
(A) Schematic of the sampling design. Left: Overall experimenter protocol for feeding and observing fishes during a single trial. Middle: Subject design. N = 19 fishes were individually housed and fed prey one at a time using a plastic pipette during a prey presentation. Right: Trials consisted of 10 prey presentations, one for each prey type and treatment combination. Prey were: 2 positive controls, 2 types of adult, female ostracod (a bioluminescent species = highlighted in blue; a nonluminescent species = highlighted in red), and 4 treatments for the two types of ostracod (alive = bold; anesthetized = italics; frozen or boiled = underlined). During a prey presentation, each fish was given the prey and watched for 1 min. We recorded the following behaviors in 20 s intervals during that one minute: number of bites taken at the food, number of times the food was rejected or spit out, number of bioluminescent events visible (for the bioluminescent prey), and if the prey was eaten. If after 1 min. the prey was not eaten, we moved on to the next fish and left the prey in the container. This process proceeded for all fish to receive a single prey. Afterwards, we returned to any fish that had not eaten their prey and marked whether the prey was still in the container or not. If it was gone, we marked it as eaten; if it was still there, we marked it as uneaten and removed it. We then repeated this process with the next prey, until all fish had received all 10 prey. This constituted a single trial. The order of prey presentations were randomized in each trial. Each fish experienced four trials. (B) Schematic of the experimental protocol, as described above. A trial is represented by a bold box outline, and within each trial, a feeding presentation for all fish is represented by the thinner box outline. These units are repeated over time. Within each presentation, the observation time increases on the y-axis, across all fish on the x-axis. Between trials, time increases horizontally, as indicated by the arrows leading to repeated units of trials and feeding presentations.

**Figure S2.**
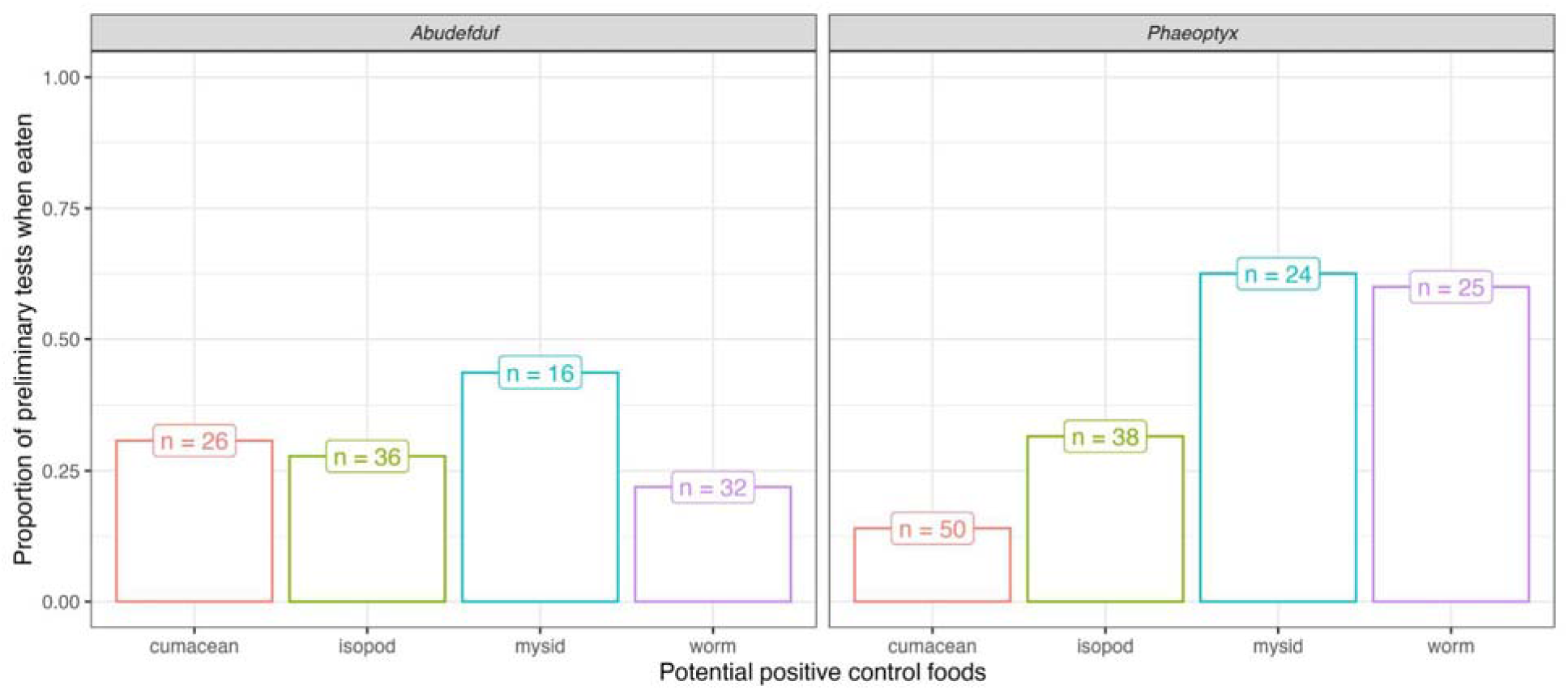
Proportion of each potential positive control prey eaten in preliminary tests when fish ate at least one prey offered. Faceted by fish genus. Isopods were chosen as they were roughly equally palatable between fish species and readily abundant compared to other potential prey.

**Figure S3.**
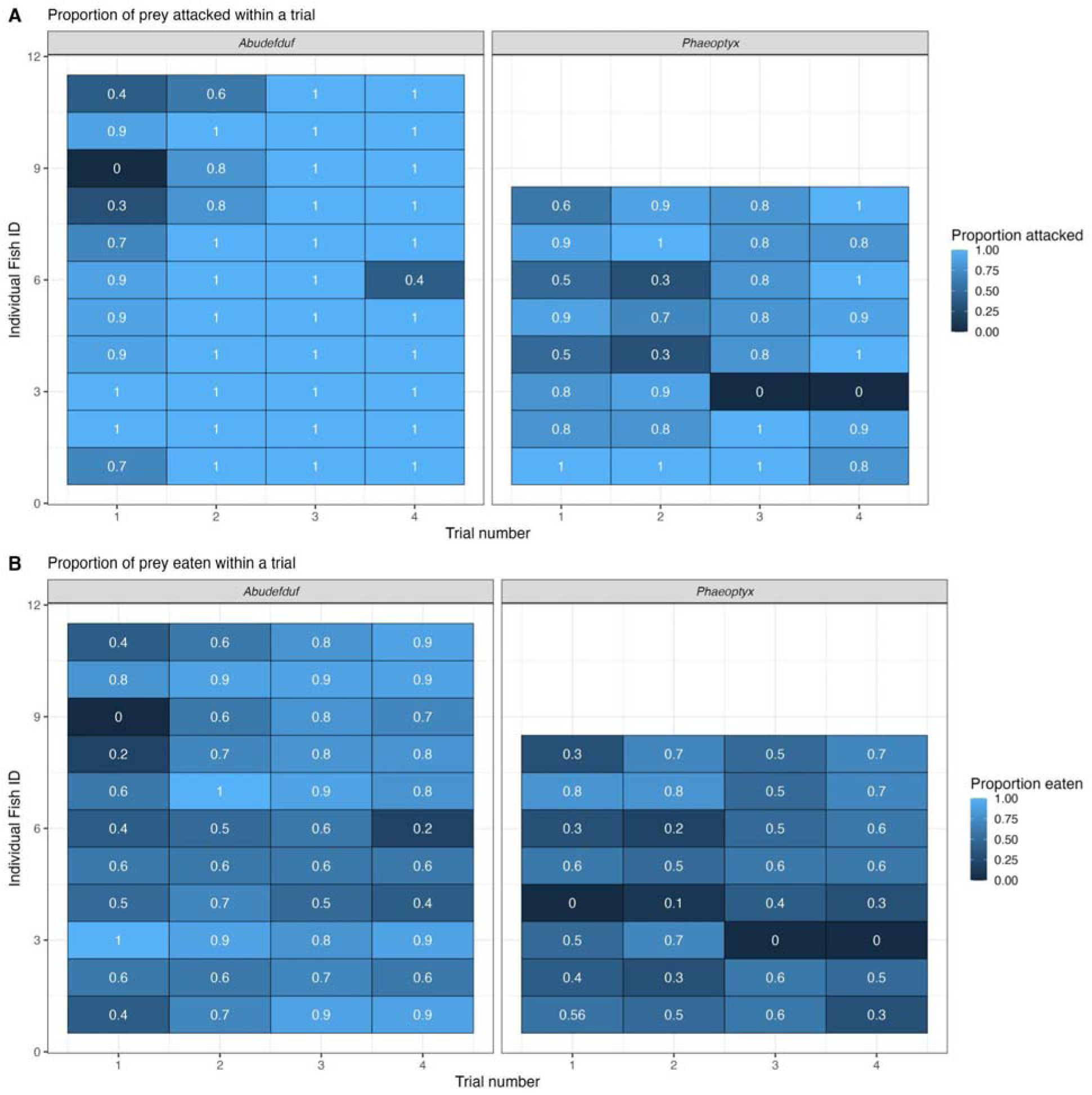
(A) Proportion of prey attacked across trials (x-axis) for each fish (y-axis). Faceted by fish genus. Individual trials where a fish failed to attack any prey item were excluded from the model assessing the risk of being eaten. (B) Proportion of prey eaten across trials (x-axis) for each fish (y-axis). Faceted by fish genus. Each trial consisted of 10 prey items.

**Figure S4.**
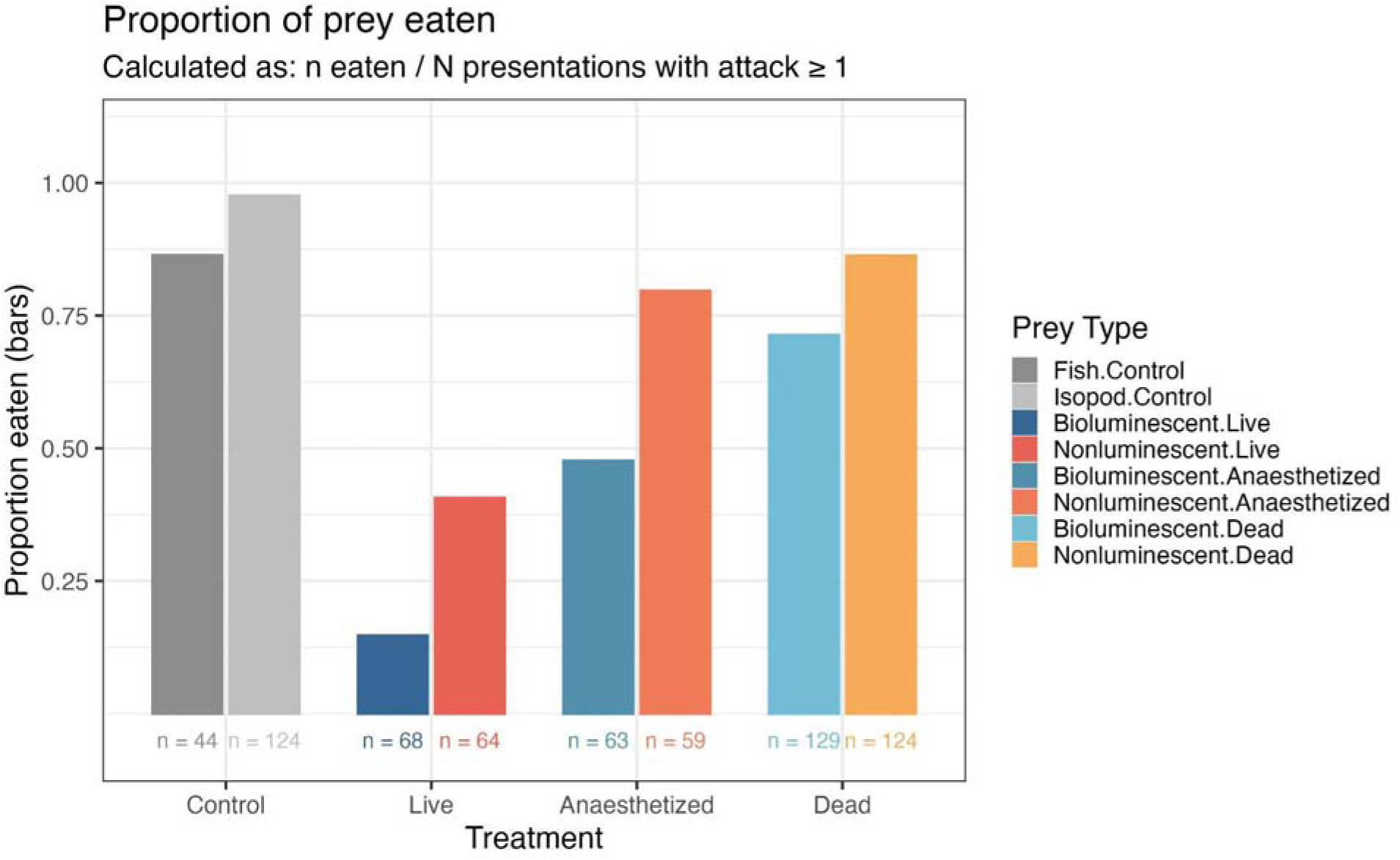
Proportion of prey eaten across all food presentations in experimental trials, including the two positive controls. Note that even for “positive” control food items, feeding proportions never reach 100% for either fish muscle or boiled cirolanid isopods.

**Figure S5.**
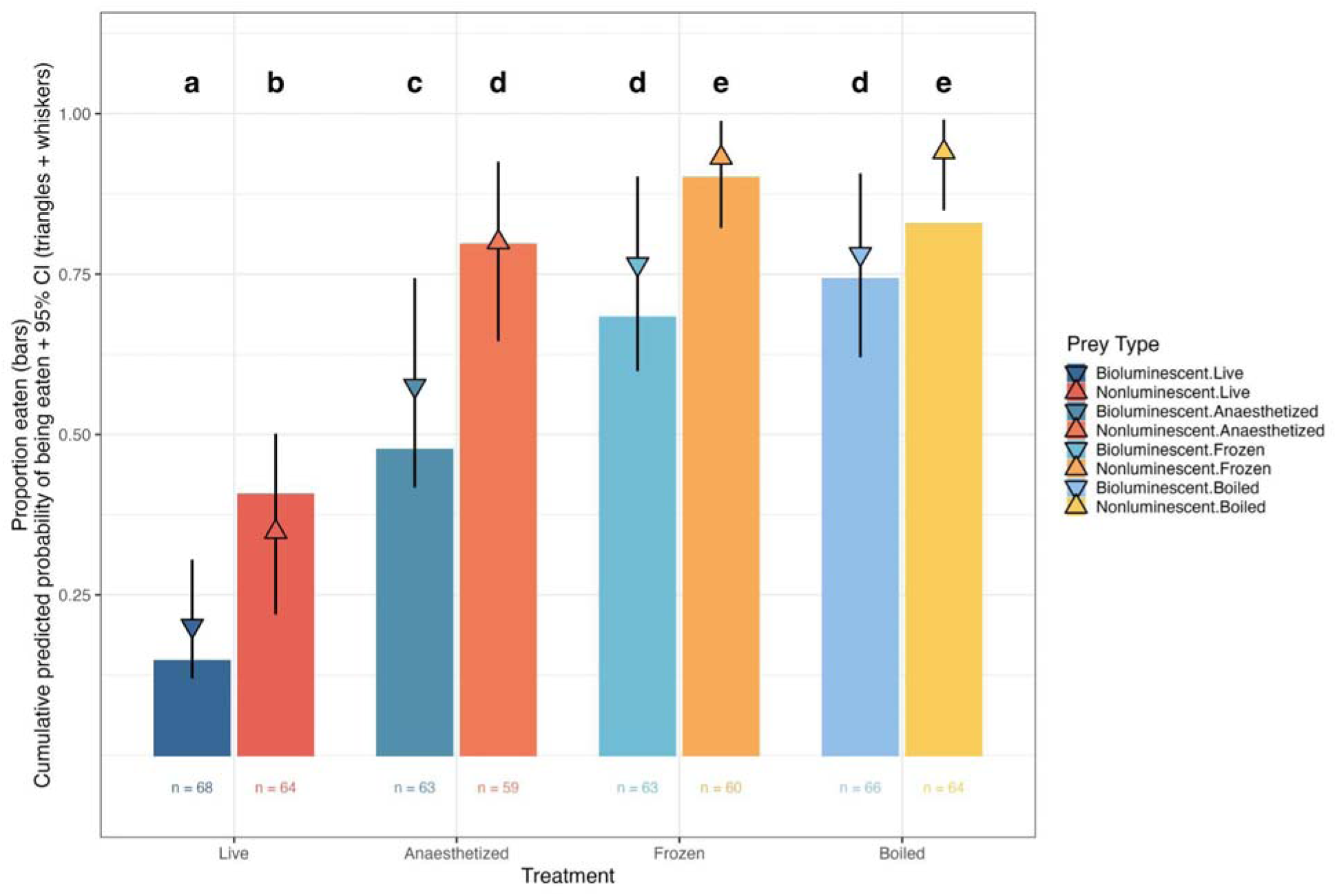
Proportion of prey eaten across presentations in experimental trials. Data plotted are the same as in Fig. 2 but with “Dead” categorized according to the original treatment levels: either Frozen or Boiled. Results are statistically indistinguishable between the Frozen and Boiled treatments, motivating us to combine them into one treatment in the main text. The conclusions remain unchanged with either categorization of the data. colours as in Fig. 2

**Figure S6.**
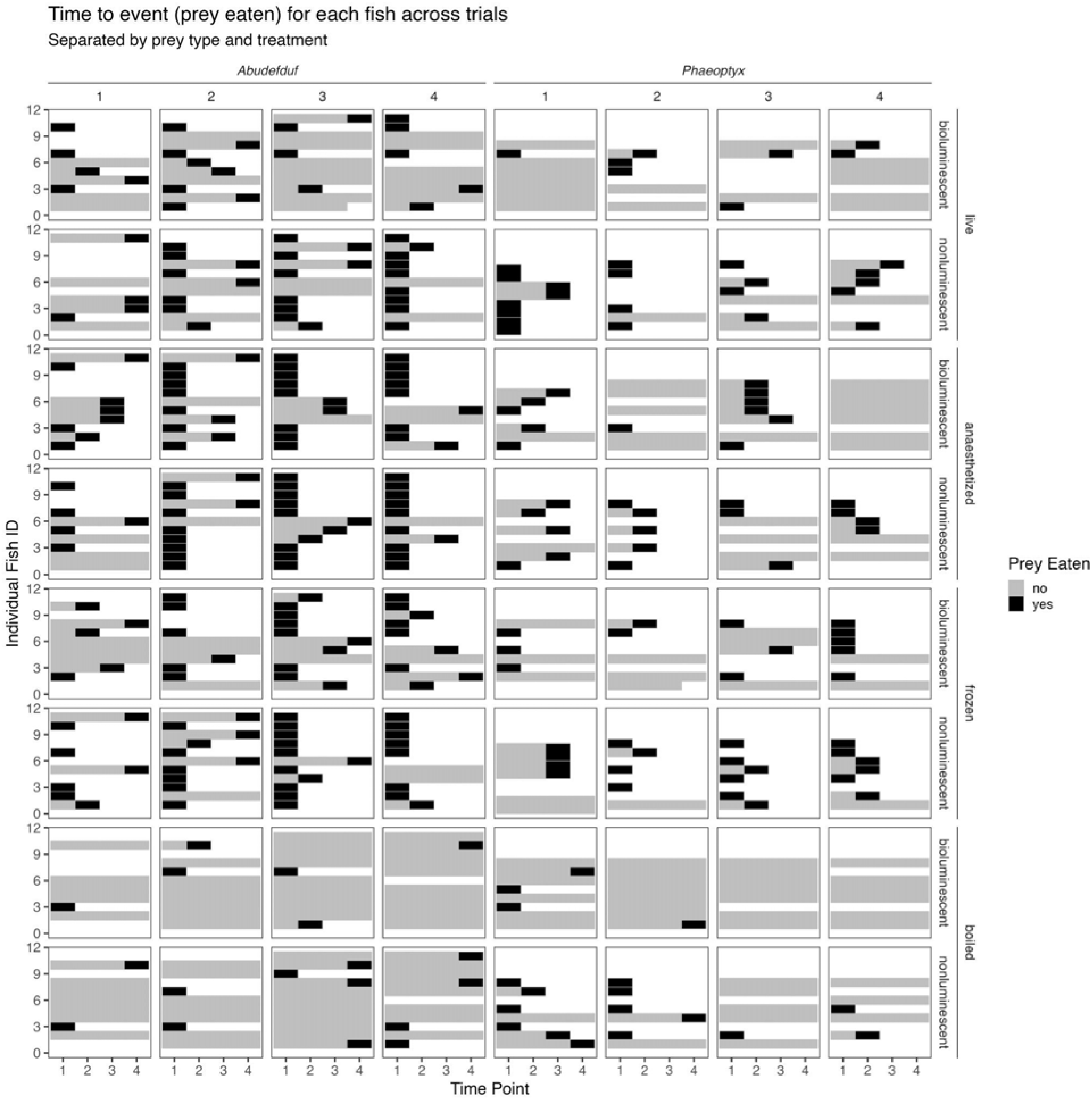
Time to event (being eaten) for every fish (y-axis) across all time points (x-axis) within each trial, across all trials (columnwise subfacets) for each predator genus (columnwise facets), for all prey treatments and types (rowwise facets and subfacets, respectively). Prey is uneaten (grey) until it is eaten (black), if at all. This is a visual representation of all the raw data, which is then filtered as described in the main body for use in the discrete-time hazard model to understand variation in the probability to be consumed.

**Figure S7.**
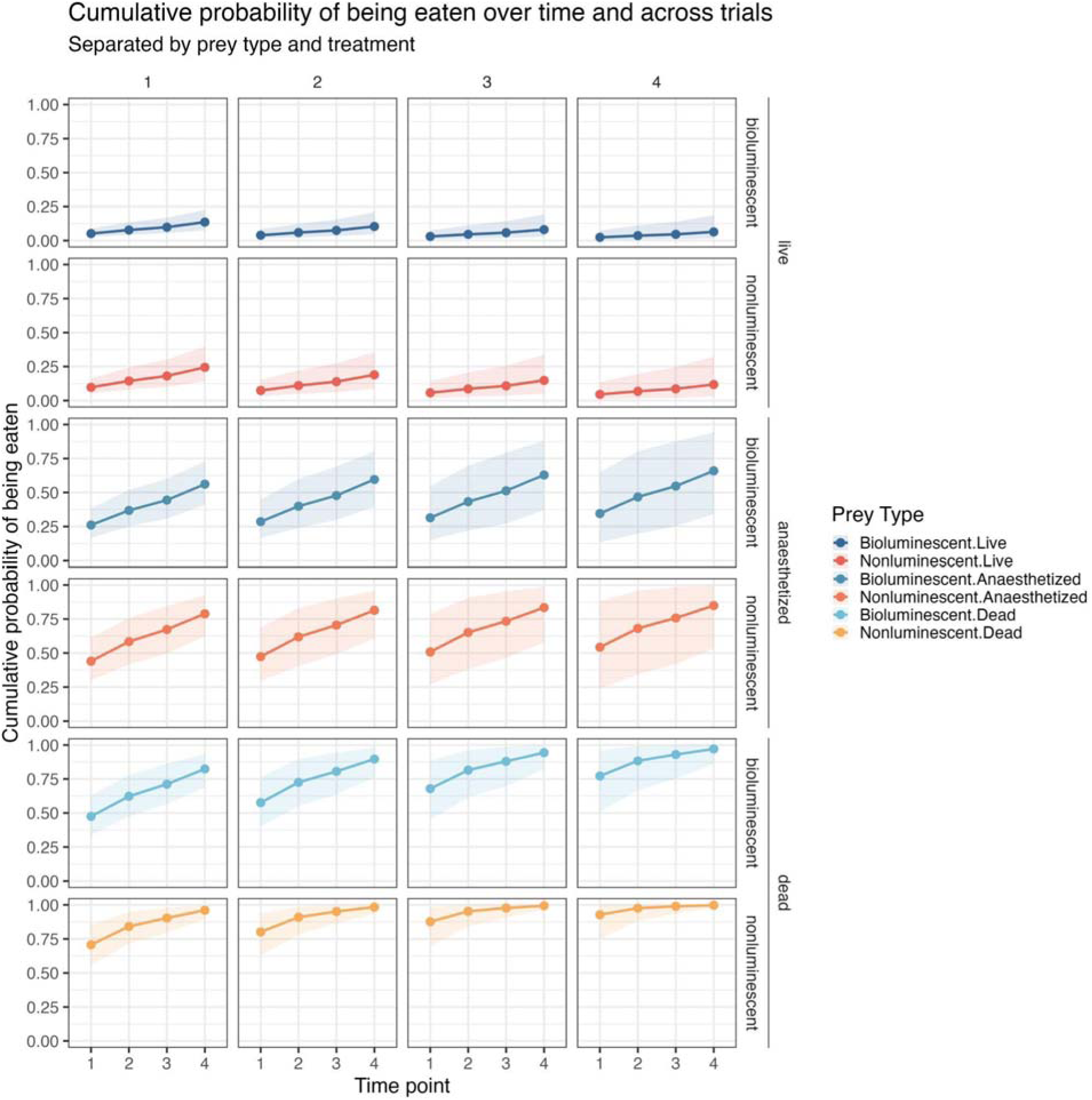
Relative probability of being eaten over time (x-axis) and across trials (column-wise facets), separated by prey type and treatment (row-wise subfacets and facets, respectively). For the plots within the main paper, the cumulative probability is the product of the relative probability calculated across all four time points; data per fixed effect are then presented with other variables held at their mean value to produce a marginal cumulative probability.

**Figure S8.**
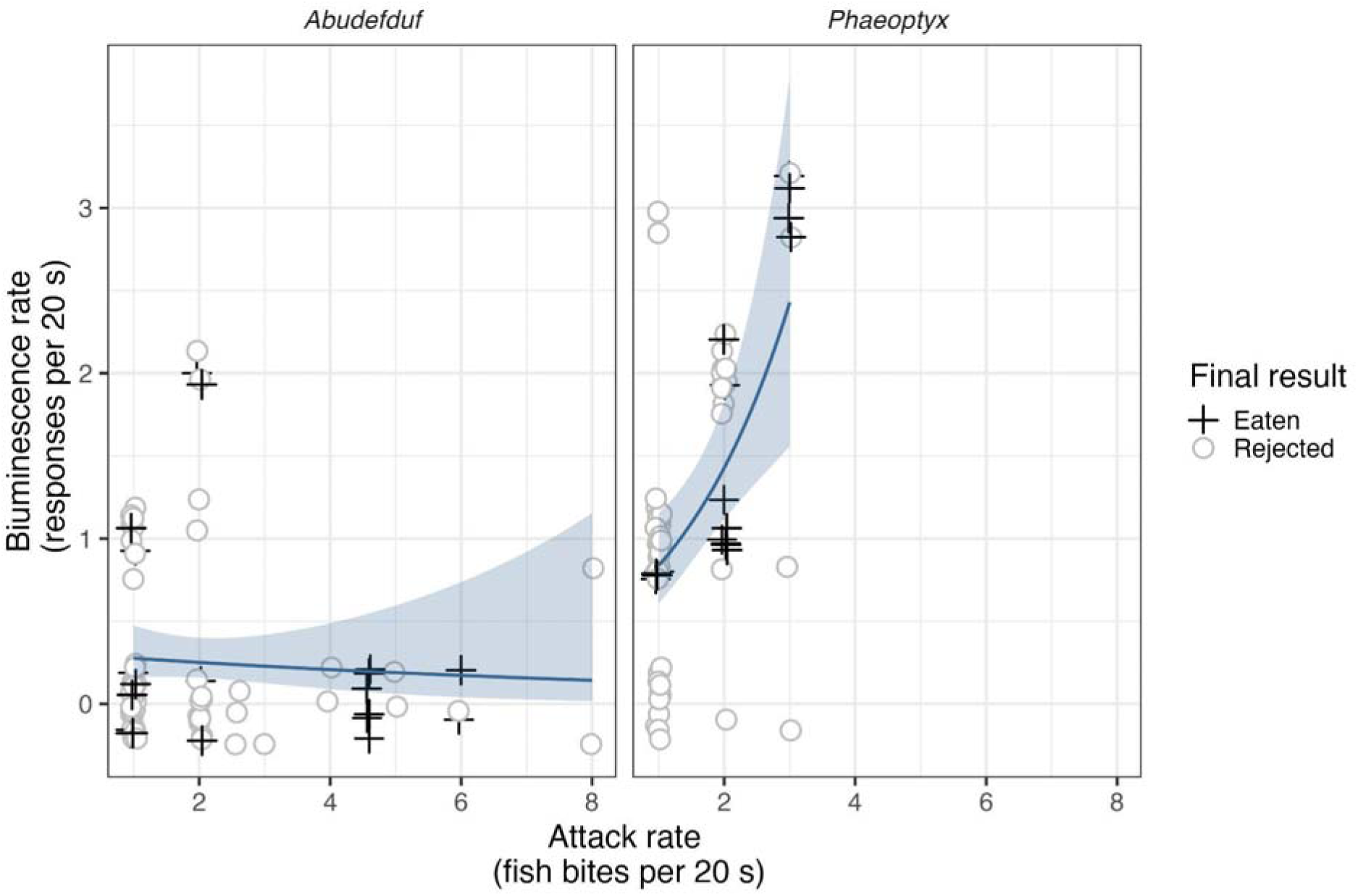
Model from Fig. 1 but with one outlier removed. Rates of bioluminescent responses increase with rates of fish attacks for *Phaeoptyx* but not *Abudefduf* predators. Results are qualitatively unchanged from the main text, even without the single outlier. See main text and Tables S1,S2 for results.

**Figure S9.**
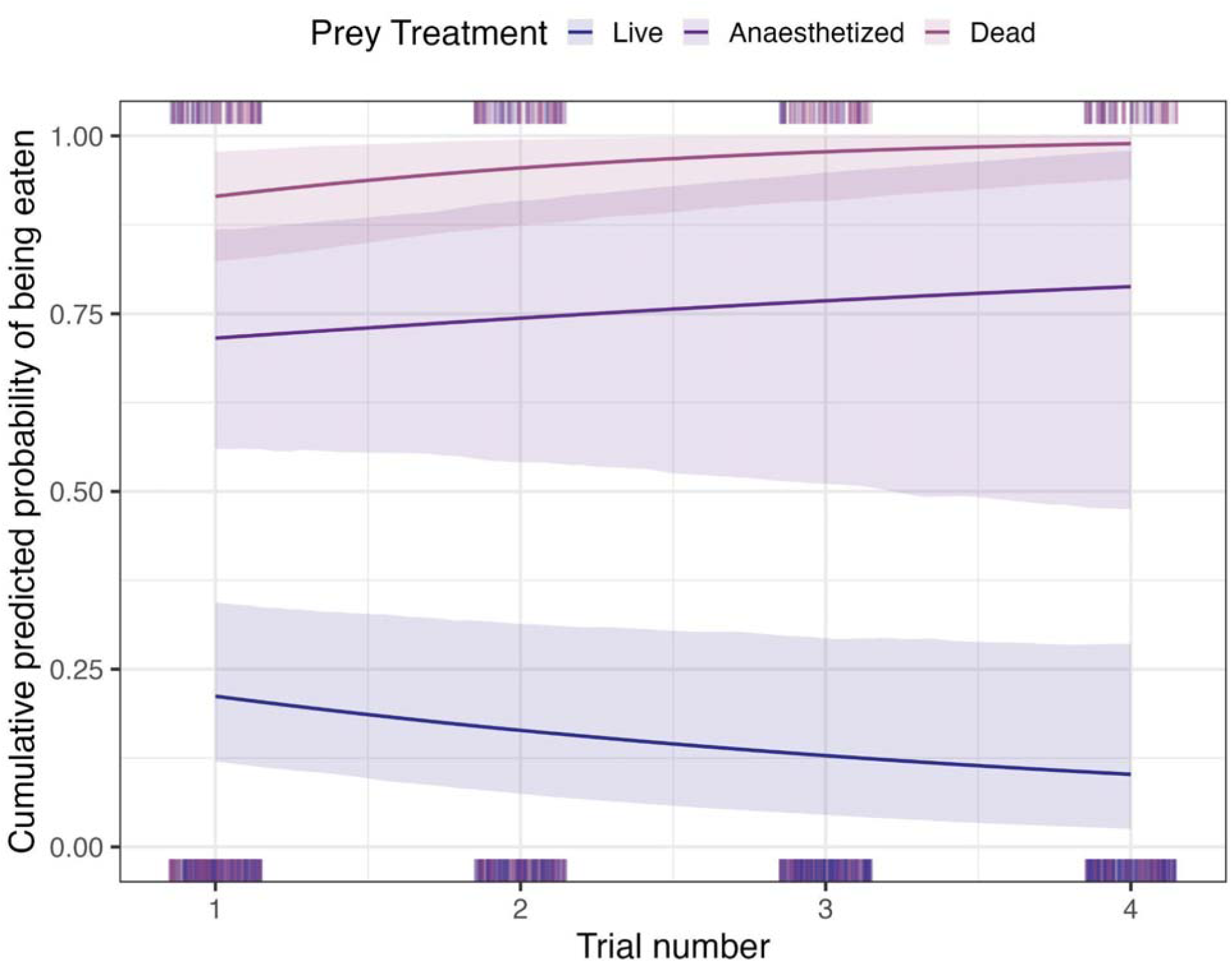
Cumulative probability of being eaten for different prey treatments as a function of trial number. Live prey had a low and decreasing cumulative probability of being eaten with more trials. Model predictions are marginal cumulative probabilities with all other fixed effects held at their mean and plotted on the original scale. Shaded regions are 95% confidence intervals from bootstrapping. Y-axis is the cumulative probability of being eaten across all four time points for a given level of bioluminescence experience, with 0 being no cumulative probability of being eaten, and 1 being 100% cumulative probability of being eaten.

**Figure S10.**
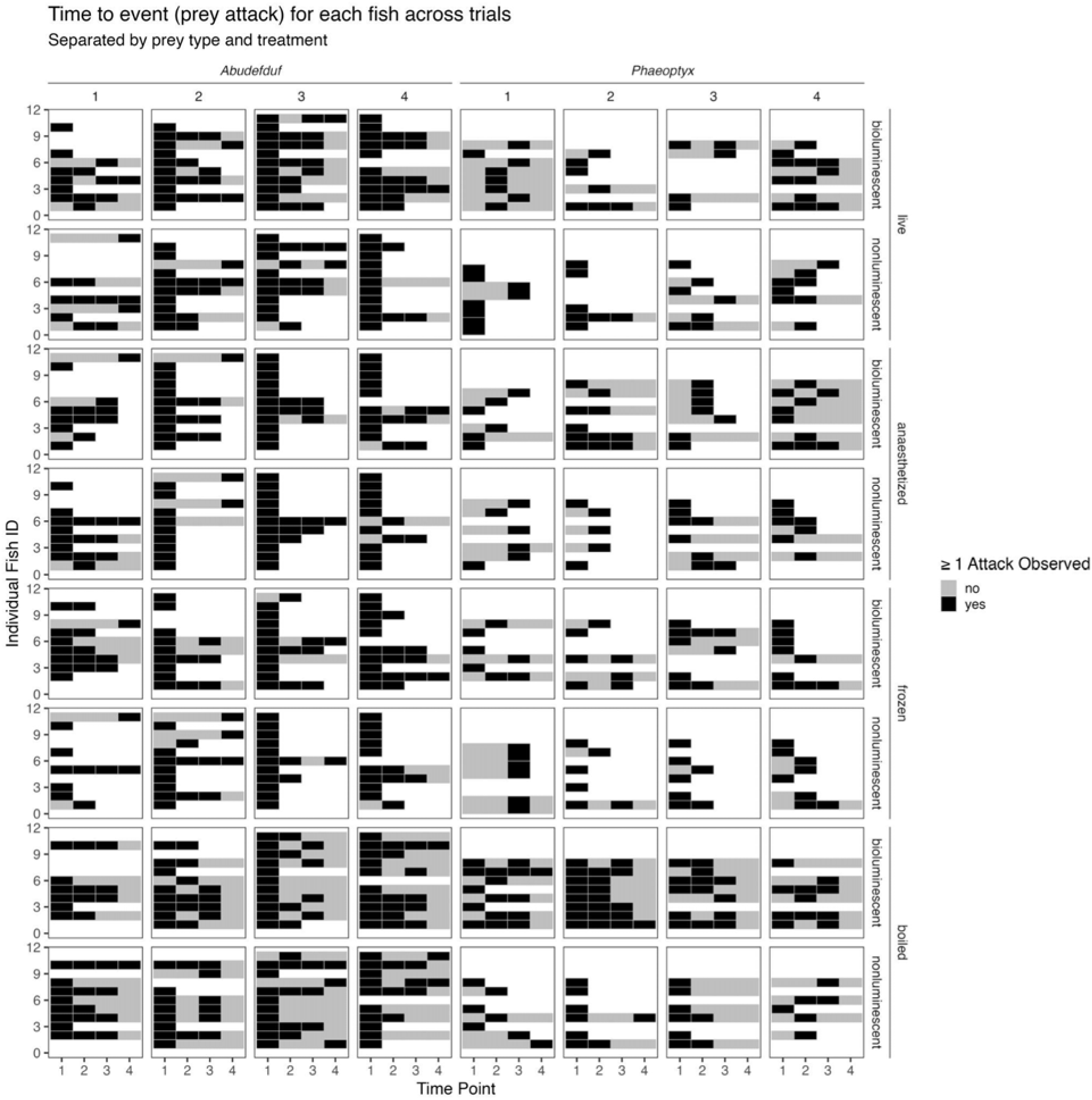
Time to event (attacking prey) for every fish (y-axis) across all time points (x-axis) within each trial, across all trials (columnwise subfacets) for each predator genus (columnwise facets), for all prey treatments and types (rowwise facets and subfacets, respectively). Prey is unmolested (grey) until it is attacked (black), if at all. Unlike in Fig. S4 where prey is eaten, prey can transition from being attacked, then rejected and left alone, to being attacked again within a presentation. This is a visual representation of all the raw data, which is then filtered as described in the supplement for use in the discrete-time hazard model to understand variation in the probability to attack.

**Figure S11.**
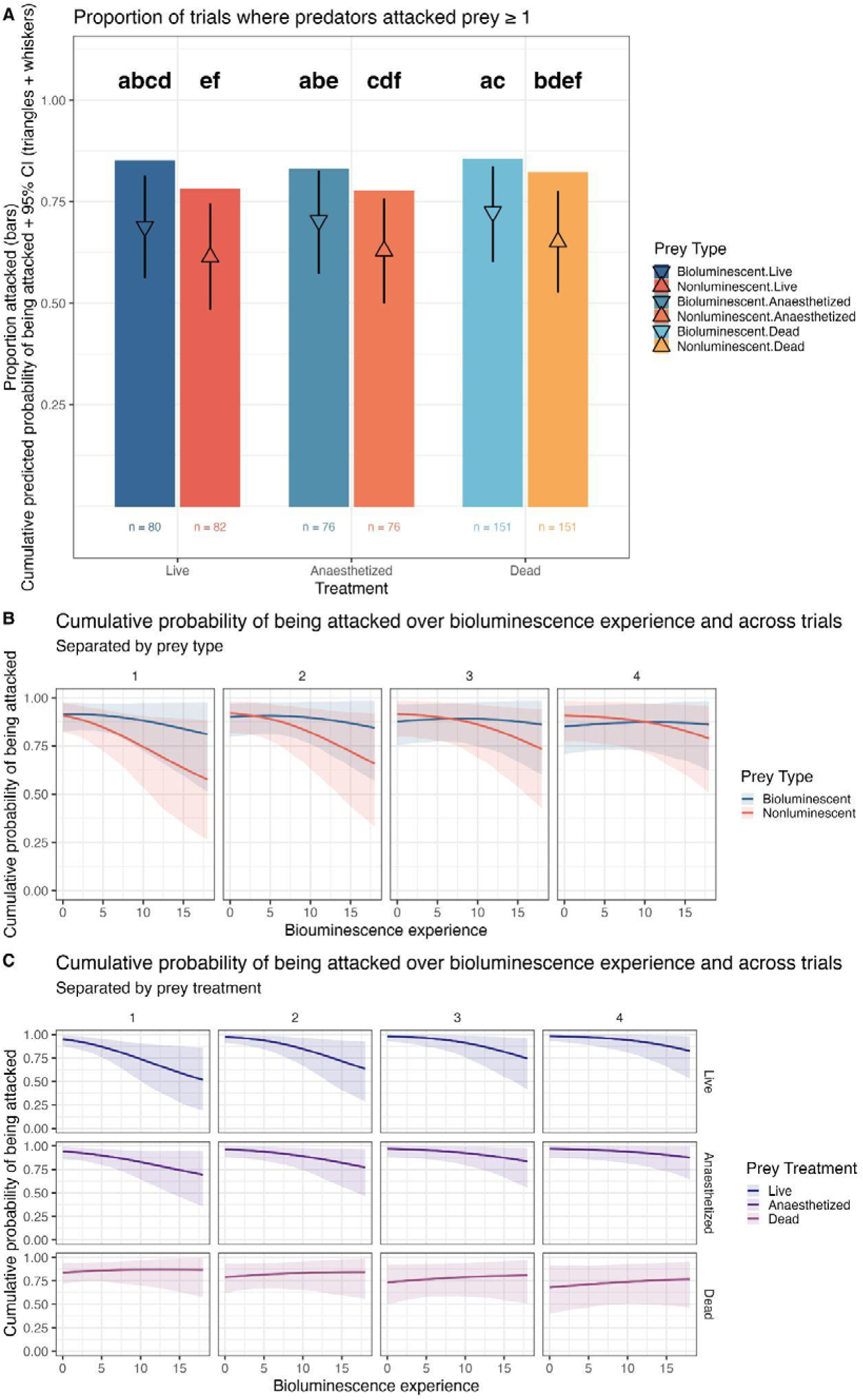
Predicted discrete-time hazard model effects showing the marginal cumulative probability of being attacked by (A) prey type and treatment (colours as in Fig. 1); (B) prey type (colours; blue = bioluminescent, red = nonluminescent) type across trials (facets) and bioluminescence experience (x-axis); and (C) prey treatment (colours; dark purple = live, light purple = anaesthetized, bright purple = dead) across trials (facets) and bioluminescence experience (x-axis).

**Figure S12.**
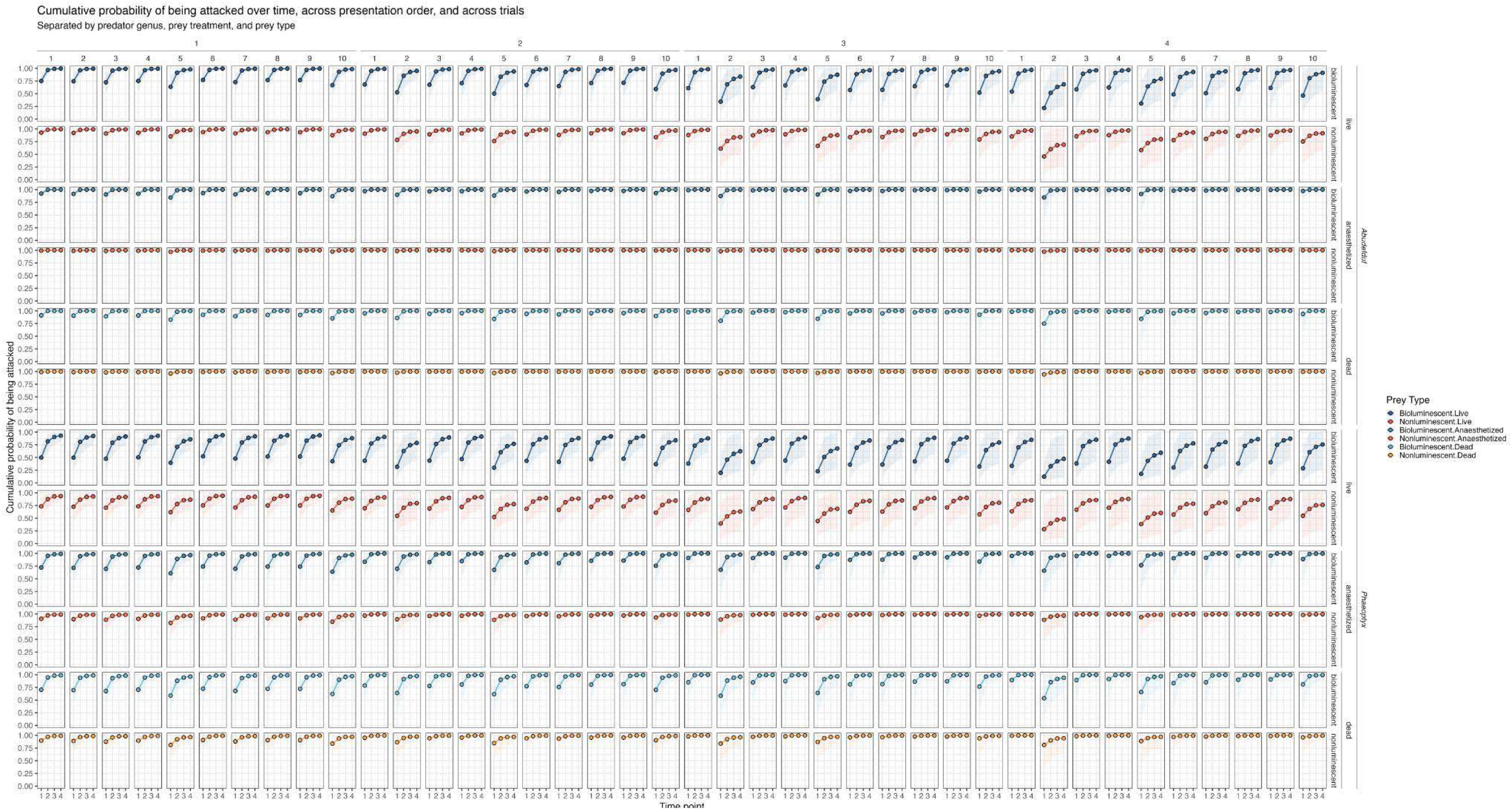
Relative probability of being attacked over time (x-axis) and across presentations and trials (column-wise sub-facets and facets, respectively), separated by prey type and treatment, and predator genus (row-wise minor facets, subfacets, and facets, respectively). For the plots in Fig. S12, the cumulative probability is the product of the relative probability calculated across all four time points; data per fixed effect are then presented with other variables held at their mean value to produce a marginal cumulative probability.

## References

1. Drinkwater E et al. 2022 A synthesis of deimatic behaviour. Biol. Rev. Camb. Philos. Soc. 97, 2237–2267. (doi:10.1111/brv.12891)

2. Umbers KDL, De Bona S, White TE, Lehtonen J, Mappes J, Endler JA. 2017 Deimatism: a neglected component of antipredator defence. Biol. Lett. 13, 20160936. (doi:10.1098/rsbl.2016.0936)

3. Umbers KDL, Lehtonen J, Mappes J. 2015 Deimatic displays. Curr. Biol. 25, R58–9. (doi:10.1016/j.cub.2014.11.011)

4. Stevens M, Ruxton GD. 2012 Linking the evolution and form of warning coloration in nature. Proc. Biol. Sci. 279, 417–426. (doi:10.1098/rspb.2011.1932)

5. Barnett JB, Scott-Samuel NE, Cuthill IC. 2016 Aposematism: balancing salience and camouflage. Biol. Lett. 12, 20160335. (doi:10.1098/rsbl.2016.0335)

6. Rojas B, Valkonen J, Nokelainen O. 2015 Aposematism. Curr. Biol. 25, R350–1. (doi:10.1016/j.cub.2015.02.015)

7. White TE, Umbers KDL. 2021 Meta-analytic evidence for quantitative honesty in aposematic signals. Proc. Biol. Sci. 288, 20210679. (doi:10.1098/rspb.2021.0679)

8. Maan ME, Cummings ME. 2012 Poison frog colors are honest signals of toxicity, particularly for bird predators. Am. Nat. 179, E1–14. (doi:10.1086/663197)

9. Summers K, Clough ME. 2001 The evolution of coloration and toxicity in the poison frog family (Dendrobatidae). Proc. Natl. Acad. Sci. U. S. A. 98, 6227–6232. (doi:10.1073/pnas.101134898)

10. Bezzerides AL, McGraw KJ, Parker RS, Husseini J. 2007 Elytra color as a signal of chemical defense in the Asian ladybird beetle Harmonia axyridis. Behav. Ecol. Sociobiol. 61, 1401–1408. (doi:10.1007/s00265-007-0371-9)

11. van den Berg CP, Santon M, Endler JA, Drummond L, Dawson BR, Santiago C, Weber N, Cheney KL. 2024 Chemical defences indicate bold colour patterns with reduced variability in aposematic nudibranchs. Proc. Biol. Sci. 291, 20240953. (doi:10.1098/rspb.2024.0953)

12. Cortesi F, Cheney KL. 2010 Conspicuousness is correlated with toxicity in marine opisthobranchs: Aposematic signals in opisthobranchs. J. Evol. Biol. 23, 1509–1518. (doi:10.1111/j.1420-9101.2010.02018.x)

13. Sherratt TN, Beatty CD. 2003 The evolution of warning signals as reliable indicators of prey defense. Am. Nat. 162, 377–389. (doi:10.1086/378047)

14. Kikuchi DW, Herberstein ME, Barfield M, Holt RD, Mappes J. 2021 Why aren’t warning signals everywhere? On the prevalence of aposematism and mimicry in communities. Biol. Rev. Camb. Philos. Soc. 96, 2446–2460. (doi:10.1111/brv.12760)

15. Chouteau M, Arias M, Joron M. 2016 Warning signals are under positive frequency-dependent selection in nature. Proc. Natl. Acad. Sci. U. S. A. 113, 2164–2169. (doi:10.1073/pnas.1519216113)

16. Endler JA. 1988 Frequency-dependent predation, crypsis and aposematic coloration. Philos. Trans. R. Soc. Lond. B Biol. Sci. 319, 505–523. (doi:10.1098/rstb.1988.0062)

17. Mappes J, Marples N, Endler JA. 2005 The complex business of survival by aposematism. Trends Ecol. Evol. 20, 598–603. (doi:10.1016/j.tree.2005.07.011)

18. Gittleman JL, Harvey PH. 1980 Why are distasteful prey not cryptic? Nature 286, 149–150. (doi:10.1038/286149a0)

19. Loeffer-Henry K, Kang C, Sherratt TN. 2025 Hidden colour signals as key drivers in the evolution of anti-predator coloration and defensive behaviours in snakes. Nat. Commun. 17, 1050. (doi:10.1038/s41467-025-67809-y)

20. Loeffler-Henry K, Kang C, Sherratt TN. 2023 Evolutionary transitions from camouflage to aposematism: Hidden signals play a pivotal role. Science 379, 1136–1140. (doi:10.1126/science.ade5156)

21. Song W, Lee S-I, Jablonski PG. 2020 Evolution of switchable aposematism: insights from individual-based simulations. PeerJ 8, e8915. (doi:10.7717/peerj.8915)

22. Kang C, Cho H-J, Lee S-I, Jablonski PG. 2016 Post-attack aposematic display in prey facilitates predator avoidance learning. Front. Ecol. Evol. 4. (doi:10.3389/fevo.2016.00035)

23. O’Hanlon JC, Rathnayake DN, Barry KL, Umbers KDL. 2018 Post-attack defensive displays in three praying mantis species. Behav. Ecol. Sociobiol. 72. (doi:10.1007/s00265-018-2591-6)

24. Higginson AD, Ruxton GD. 2010 Optimal defensive coloration strategies during the growth period of prey. Evolution 64, 53–67. (doi:10.1111/j.1558-5646.2009.00813.x)

25. Lau ES, Oakley TH. 2021 Multi-level convergence of complex traits and the evolution of bioluminescence. Biol. Rev. Camb. Philos. Soc. 96, 673–691. (doi:10.1111/brv.12672)

26. Claes JM, Haddock SHD, Coubris C, Mallefet J. 2024 Systematic distribution of bioluminescence in marine animals: A species-level inventory. Life (Basel) 14, 432. (doi:10.3390/life14040432)

27. Haddock SHD, Moline MA, Case JF. 2010 Bioluminescence in the sea. Ann. Rev. Mar. Sci. 2, 443–493. (doi:10.1146/annurev-marine-120308-081028)

28. Underwood TJ, Tallamy DW, Pesek JD. 1997 Bioluminescence in firefly larvae: a test of the aposematic display hypothesis (Coleoptera: Lampyridae). J. Insect Behav. 10, 365–370. (doi:10.1007/bf02765604)

29. De Cock R. 2003 Glow-worm larvae bioluminescence (Coleoptera: Lampyridae) operates as an aposematic signal upon toads (Bufo bufo). Behav. Ecol. 14, 103–108. (doi:10.1093/beheco/14.1.103)

30. Cock RD, Matthysen E. 1999 Aposematism and Bioluminescence: Experimental evidence from Glow-worm Larvae(Coleoptera: Lampyridae). Evol. Ecol. 13, 619–639. (doi:10.1023/a:1011090017949)

31. Marek P, Papaj D, Yeager J, Molina S, Moore W. 2011 Bioluminescent aposematism in millipedes. Curr. Biol. 21, R680–1. (doi:10.1016/j.cub.2011.08.012)

32. Long SM, Lewis S, Jean-Louis L, Ramos G, Richmond J, Jakob EM. 2012 Firefly flashing and jumping spider predation. Anim. Behav. 83, 81–86. (doi:10.1016/j.anbehav.2011.10.008)

33. Livermore J, Perreault T, Rivers T. 2018 Luminescent defensive behaviors of polynoid polychaete worms to natural predators. Mar. Biol. 165. (doi:10.1007/s00227-018-3403-2)

34. Cohen AC, Morin JG. 2003 Sexual morphology, reproduction and the evolution of bioluminescence in Ostracoda. Paleontol. Soc. Pap. 9, 37–70. (doi:10.1017/s108933260000214x)

35. Rivers TJ, Morin JG. 2008 Complex sexual courtship displays by luminescent male marine ostracods. J. Exp. Biol. 211, 2252–2262. (doi:10.1242/jeb.011130)

36. Duchatelet L, Dupont S. 2025 Marine eukaryote bioluminescence: a review of species and their functional biology. Mar. Life Sci. Technol. 7, 366–381. (doi:10.1007/s42995-024-00250-0)

37. Letendre F, Twardowski M, Blackburn A, Poulin C, Latz MI. 2024 A review of mechanically stimulated bioluminescence of marine plankton and its applications. Front. Mar. Sci. 10. (doi:10.3389/fmars.2023.1299602)

38. Grober MS. 1988 Brittle-star bioluminescence functions as an aposematic signal to deter crustacean predators. Anim. Behav. 36, 493–501. (doi:10.1016/s0003-3472(88)80020-4)

39. Hanley KA, Widder EA. 2017 Bioluminescence in Dinoflagellates: Evidence that the Adaptive Value of Bioluminescence in Dinoflagellates is Concentration Dependent. Photochem. Photobiol. 93, 519–530. (doi:10.1111/php.12713)

40. Rivers TJ, Morin JG. 2012 The relative cost of using luminescence for sex and defense: light budgets in cypridinid ostracods. J. Exp. Biol. 215, 2860–2868. (doi:10.1242/jeb.072017)

41. Ellis EA et al. 2023 Sexual signals persist over deep time: Ancient co-option of bioluminescence for courtship displays in cypridinid ostracods. Syst. Biol. 72, 264–274. (doi:10.1093/sysbio/syac057)

42. Abe K, Ono T, Yamada K, Yamamura N, Ikuta K. 2000 Multifunctions of the upper lip and a ventral reflecting organ in a bioluminescent ostracod Vargula hilgendorfii (Müller, 1890). In Evolutionary Biology and Ecology of Ostracoda, pp. 73–82. Dordrecht: Springer Netherlands. (doi:10.1007/978-94-017-1508-9_5)

43. Huvard AL. 1993 Ultrastructure of the light organ and immunocytochemical localization of luciferase in luminescent marine ostracods (Crustacea: Ostracoda: Cypridinidae). J. Morphol. 218, 181–193. (doi:10.1002/jmor.1052180207)

44. Hensley NM et al. 2021 Selection, drift, and constraint in cypridinid luciferases and the diversification of bioluminescent signals in sea fireflies. Mol. Ecol. 30, 1864–1879. (doi:10.1111/mec.15673)

45. Tsarkova AS. 2021 Luciferins under construction: A review of known biosynthetic pathways. Front. Ecol. Evol. 9. (doi:10.3389/fevo.2021.667829)

46. Torres E, Morin JG. 2007 Vargula annecohenae, a new species of bioluminescent ostracode (Myodocopida: Cypridinidae) from Belize. J. Crustacean Biol. 27, 649–659. (doi:10.1651/s-2769.1)

47. Poulsen. 1962 Ostracoda-Myodocopa. BRILL. (doi:10.1163/9789004610989)

48. Cohen AC, Oakley TH. 2017 Collecting and processing marine ostracods. J. Crustacean Biol. 37, 347–352. (doi:10.1093/jcbiol/rux027)

49. Johnson FH, Shimomura O. 1978 [32] introduction to the Cypridina system. In Bioluminescence and Chemiluminescence, pp. 331–364. Elsevier. (doi:10.1016/0076-6879(78)57034-1)

50. Clark TG, Bradburn MJ, Love SB, Altman DG. 2003 Survival analysis part I: basic concepts and first analyses. Br. J. Cancer 89, 232–238. (doi:10.1038/sj.bjc.6601118)

51. 2014 Fish fireworks: Bioluminescence. BBC, 11 August. See https://www.bbc.co.uk/programmes/p024m0s1.

52. Mesrop LY, Minsky G, Drummond MS, Goodheart JA, Proulx SR, Oakley TH. 2024 Ancient secretory pathways contributed to the evolutionary origin of an ecologically impactful bioluminescence system. bioRxiv. (doi:10.1101/2024.05.07.593075)

53. Prével A, Krebs RM. 2021 Higher-order conditioning with simultaneous and backward conditioned stimulus: Implications for models of Pavlovian conditioning. Front. Behav. Neurosci. 15, 749517. (doi:10.3389/fnbeh.2021.749517)

54. Oakley TH. 2025 Visual information in the dark: Bioluminescence and perceptual design through evolution. Funct. Ecol. 39, 2611–2625. (doi:10.1111/1365-2435.70106)

55. Burkenroad MD. 1943 A possible function of bioluminescence. J. Mar. Res. 5.

56. Buskey EJ, Swift E. 1983 Behavioral responses of the coastal copepod Acartia hudsonica (Pinhey) to simulated dinoflagellate bioluminescence. J. Exp. Mar. Bio. Ecol. 72, 43–58. (doi:10.1016/0022-0981(83)90018-7)

57. Mesinger AF, Case JF. 1992 Dinoflagellate luminescence increases susceptibility of zooplankton to teleost predation. Mar. Biol. 112, 207–210. (doi:10.1007/bf00702463)

58. Zhu C et al. 2024 Firefly toxin lucibufagins evolved after the origin of bioluminescence. PNAS Nexus 3, pgae215. (doi:10.1093/pnasnexus/pgae215)

59. Prevett A, Lindström J, Xu J, Karlson B, Selander E. 2019 Grazer-induced bioluminescence gives dinoflagellates a competitive edge. Curr. Biol. 29, R564–R565. (doi:10.1016/j.cub.2019.05.019)

60. Malcolm SB. 1986 Aposematism in a soft-bodied insect: a case for kin selection. Behav. Ecol. Sociobiol. 18, 387–393. (doi:10.1007/bf00299669)

61. Brodie ED 3rd, Agrawal AF. 2001 Maternal effects and the evolution of aposematic signals. Proc. Natl. Acad. Sci. U. S. A. 98, 7884–7887. (doi:10.1073/pnas.141075998)

62. Järvi T, Sillén-Tullberg B, Wiklund C, Jarvi T, Sillen-Tullberg B. 1981 Individual versus kin selection for aposematic coloration: A reply to Harvey and Paxton. Oikos 37, 393. (doi:10.2307/3544136)

63. Oakley TH, Halvonik-Sánchez A, Speiser DI, Hensley NM. 2026 Diel metabolic variation and the energetic demands of courtship in bioluminescent Photeros ostracods. bioRxiv. (doi:10.64898/2026.06.25.734310)

